# Flavonols contrary affect the interconnected glucosinolate and camalexin biosynthesis pathway in *Arabidopsis thaliana*

**DOI:** 10.1101/2022.10.01.510434

**Authors:** Jogindra Naik, Shivi Tyagi, Ruchika Rajput, Pawan Kumar, Boas Pucker, Naveen C. Bisht, Prashant Misra, Ralf Stracke, Ashutosh Pandey

## Abstract

Flavonols are structurally and functionally diverse molecules playing roles in plant biotic and abiotic stress tolerance, auxin transport inhibition, pollen development, etc. Despite their ubiquitous occurrence in land plants and multifunctionality, the effect of perturbation of flavonol biosynthesis over global gene expression and pathways other than flavonoid biosynthesis has not been studied in detail. To understand the signaling role of different flavonol metabolites, herein, we used the flavonol deficient *Arabidopsis thaliana* loss-of-function mutant *flavonol synthase1 (fls1-3)* as object of study. Comparative transcriptome and metabolic profiling were used to study the effects of genetic flavonol deficiency and exogenous supplementation with flavonol derivatives (kaempferol, quercetin and rutin) on different cellular processes in the seedling. Various flavonol biosynthesis-related regulatory and structural genes were found to be up-regulated in the *fls1-3* mutant which could be reversed by exogenous flavonol feeding. Our manifold comparative studies indicated the modulation of various biological processes and metabolic pathways by flavonols. Camalexin biosynthesis was found to be negatively regulated by flavonols. Interestingly, flavonols appeared to promote the accumulation of aliphatic glucosinolate through transcription factor-mediated up-regulation of biosynthesis genes. Overall, this study provides new insights into molecular mechanisms by which flavonols interfere with the relevant signal chains and their molecular targets and adds new knowledge to the expanding plethora of biological activity of flavonols in plants.

**Significance:** Comparative transcriptome and metabolomic profiling of genetic flavonol deficiency and exogenous flavonol supplementation in *A. thaliana* seedlings, for the first-time revealed the inverse regulation of interconnected specialized metabolite pathways by flavonol aglycones, and -glycosides. Flavonols negatively regulate camalexin biosynthesis, while promoting the accumulation of aliphatic glucosinolates. Our study adds new insights into the expanding plethora of biological activity of flavonols in plants and will help to uncover the molecular mechanisms by which flavonols interfere with the relevant signal chains and their molecular targets.

## Introduction

Flavonoids are considered as a group of plant specialized metabolites of ubiquitous occurrence (1, 2). These compounds have been reported to play roles in stress response, ROS (reactive oxygen species) mediated plant developmental process, and also act as signaling molecule in the regulation of gene expression (3–5). Recently, flavonoids have been connected to the circadian cycling in plants (6). Flavonols, a sub-class of flavonoids, have also been implicated in different developmental processes such as root growth, lateral root development, pollen tube growth, stomatal movement, etc., besides their classical role in biotic and abiotic stress resistance (7–12). For obvious reasons, flavonols are apparently more integral component of plant signaling machinery than just existing as “secondary metabolites”. The flavonol biosynthesis branches emerges early from the flavonoid pathway with the flavonol synthase (FLS1) as first committed rate-limiting enzyme (13, 14). At transcription level, flavonol biosynthesis is mainly regulated by SG7 R2R3 MYB transcription factors MYB12, MYB111, MYB11 independent of other copartners in Arabidopsis (15–17). Besides these regulators, AtMYB21, AtMYB24, and AtMYB57 are also known to regulate the expression of *AtFLS1* in inflorescence and flowering tissues in Arabidopsis (18, 19).

Flavonoids are biosynthesized in various tissues under specific conditions and are capable to move from cell to cell (5, 20, 21). The differential movement of flavonoids can be attributed to their structural diversity. For example, naringenin, dihydrokaempferol and dihydroquercetin, when applied exogenously, were absorbed at root tips, middle portions, and cotyledons of seedlings, and, thereafter moved to distal tissues. Later, they were converted to quercetin and kaempferol. On the contrary, kaempferol and quercetin are absorbed at the root tips only and could not move upwards (20). Experiments have shown that uptake of exogenous flavonoids is an active transport, and mediated by ABC transporters. Intracellular and inter-cellular transport of endogenous flavonoids have also been reported, which has been shown to be mediated by ABCC, P-ATPase and MATE families of transporters (20, 22, 23). Glutathione S-transferase like proteins called ligandin has been recently identified to protect flavonoids during vacuolar transport (24).

Several flavonoid responsive genes have been identified in Arabidopsis after exogenous supplementation of naringenin to WT and *tt5* mutants (25). Naringenin treatment led to transcriptome-wide changes involving genes responsible for cellular trafficking, stress response, and signaling. It was also observed that flavonoids down regulate expression of jasmonic acid biosynthetic genes like *LOX3*, *OPR3*, etc. (25). Quercetin treatment to the rice root brought changes in genome-wide gene expression particularly related to hormone and oxidative response, glutathione metabolism, phenylpropanoids synthesis, and hormone signal transduction (26). In Sorghum, exogenous supplementation of flavonoids glycone and aglycone negatively and positively regulated root growth and lateral roots, respectively (27). Flavonols have also been shown to be essential for pollen germination in plants. To this end, kaempferol supplementation to Petunia (*Petunia hybrida*), resulted in increased pollen germination and tube outgrowth (28). Several genes responsive to kaempferol including those involved in plant cell signalling, for example, S/D4, a leucine-rich repeat protein, S/D1, a LIM-domain protein, D14, a zinc finger protein, along with S/D10, S/D3, etc. were identified (28). Scutellarin having 6-hydroxyl and 7-glucoside moiety, has been reported to promote root length growth. This molecule was found to activate the expression of transcription factor *NUTCRACKER (NUC)* genes and promote cell division at cortex/endodermis initials. It was also observed that root growth and meristem size is negatively regulated by flavonoids lacking 6-hydroxyl and 7-glucoside (29). Flavonols display ROS scavenging activity and thereby these molecules can modify ROS mediated signaling and plant processes such as pollen tube growth, ABA induced stomatal closure, lateral root emergence (7, 8, 12). ROS is also known to regulate MPK3, MPK6 (30) as well as calcium channel (31), whereas flavonol affects several kinases-phosphatases (32). In view of the above-mentioned, it can be presumed that flavonoids exhibit signaling role by inhibiting synthesis or transport of auxin, modulating activity of kinase-phosphatase modules, or by scavenging ROS. Although few studies, as detailed above, pertaining to understanding the function of flavonols have been carried out, a detailed information about their signalling role still remains obscure.

Apart from flavonoids, two additional classes of specialized metabolites, namely glucosinoates (GSL) and camalexin are biosynthesized in Arabidopsis. Structurally, GSLs have oxime-thioglucose groups linked to a side chain derived from amino acids (33, 34). GSLs are broadly classified into three categories based on their precursor molecule -: aliphatic, indolic, and aromatic GSL (34–36). Aliphatic GSLs are derived from methionine, where different biosynthetic enzymes like MAM1 (methylthioalkylmalate synthase), CYP79F1, CYP79F2, CYP83A1 catalyze the successive steps (37, 38). Tryptophan derived indolic GSLs are synthesized by the successive catalysis of CYP79B1, CYP79B3, and several other enzymes (39–41). GSLs are further modified via methoxylation, benzoylation, hydroxylation at their side chains (34).

Transcriptionally, indolic GSL biosynthesis genes are regulated by MYB regulatory triad MYB34-MYB51-MYB122, while MYB28-MYB29-MYB76 regulate aliphatic branch (38, 40, 42–44). Under pathogen attack, GSLs are converted to toxic forms like isothiocyanate by myrosinase enzymes (45, 46). Camalexin is a 3-thiazol-2-yl-indole compound belonging to the class of indole alkaloids (47, 48). It is synthesized from tryptophan, and the biosynthetic pathway shares some initial steps with indole glucosinolate pathway. The intermediate of indole GSL biosynthesis indole acetaldoxime (IAOx) is converted to camalexin by the core biosynthesis enzymes like CYP71A13, CYP71B15 (PAD3) (14, 49). In *Arabidopsis*, camalexin is widely studied for its role in defense response to necrotrophic fungus particularly *Botrytis cinerea* (50, 51). The biosynthesis of these two groups of secondary metabolites has been shown to be regulated by diverse factors (34, 52, 53). However, cross-talk of GSL and camalexin biosynthesis with flavonols remains unexplored.

Against this backdrop, in the present study, we have utilized both flavonol deficient *fls1-3* mutant and wild type (WT) Col-0 plants to study flavonol uptake and modulation in the global gene expression following flavonol treatment. Targeted metabolite analysis revealed that flavonols are taken up and later get modified into other derivatives. Expression profiling of flavonol biosynthetic and regulatory genes after chemical complementation in mutant and wild-type revealed the possible feedback regulation of flavonol biosynthesis by exogenous flavonol supply. Further, the RNA-Seq data showed that all three metabolites brought about largescale changes in the expression level of genes related to phenylpropanoids metabolism, glucosinolate synthesis, hormone synthesis, signal transduction, root development, ROS signaling, etc. In addition to expression analysis, the quantification of camalexin and major aliphatic glucosinolates in response to different flavonol metabolites revealed that flavonols negatively regulate the camalexin biosynthesis, while promote accumulation of aliphatic glucosinolate.

## Materials and methods

### Plant materials and growth conditions

The *A. thaliana fls1-3* (Q71Stop nonsense) mutant in the Col-0 background has been described in Kuhn *et al.* (54). The seeds were fume sterilized in air tight desiccator bottle with 2 ml HCl and 80 ml NaOCl. Seeds were sown and germinated on half-strength MS media (pH 5.8) supplemented with 0.5 % sucrose and 0.8 % plant agar, but devoid of vitamins. Seeds were stratified for 48h at 4°C in the dark, before shifting to light. Plants were grown in a growth chamber with 16h light/8h dark cycles at 22°C.

### Flavonol derivative feeding

After 7 days, seedlings were transferred with tweezers to MS media supplemented with 100 μM each kaempferol, quercetin, and rutin (dissolved in ethanol). Pure ethanol was used for mock treatment. Growing was continued as described above. Seedlings were harvested after 24h, washed 2 times with distilled water followed by soaking excessive water in tissue wipe, and frozen in liquid nitrogen and stored at −80°C.

### Quantitative real-time RT-PCR (RT-qPCR)

Total RNA was isolated from frozen seedlings using Spectrum™ Plant Total RNA Kit (Sigma-Aldrich) followed by RNase-free DNase treatment (TURBO DNA-freeTM Kit-Invitrogen). Total RNA was further used to synthesize first-strand cDNA using RevertAid H Minus First Strand cDNA synthesis Kit (ThermoFisher) according to the manufacturer’s protocol. RT-qPCR analysis were performed with 2X SYBR Green PCR Master mix (Applied Biosystems) on a 7500 Fast Real time PCR System (Applied Biosystems) as described previously (55). Expression level of the analysed genes is given as relative fold change (2^−ΔΔCT^) in biological replicates (n=3), related to the sample having the lowest expression or the control sample, depending on the requirement. *ACTIN* (AT5G59370) expression was taken as reference. Primers used for RT-qPCR are listed in *SI Appendix* Table S1.

### RNA-Sequencing and data analysis

Total RNA was isolated from seedlings using Spectrum™ Plant Total RNA Kit (Sigma-Aldrich) followed by DNase treatment as described above. Total RNA was quantified using a NanoDrop spectrometer and integrity was checked using an Agilent 2100 Bioanalyzer system. The RNA-Seq paired end sequencing libraries were prepared from the quantity and quality check (QC) passed RNA samples using NEBNext® Ultra^TM^II directional RNA Library Prep Kit for Illumina (NEB) as per manufacturer’s instruction. The QC of libraries were analyzed on Agilent 4200 Tape Station and the sequencing was carried out on an Illumina NovaSeq 6000 platform. The raw reads were submitted to the SRA database under BioProject accession number PRJNA880681. Paired-end RNA-Seq reads were aligned to the *A. thaliana* reference genome sequence TAIR10 via STAR v2.5.1b in 2-pass mode (56). Read pairs were considered mapped if the alignment covers at least 90% of the length and has a similarity of at least 95%. Next, feature Counts v1.5.0-p3 (57) was applied to quantify gene expression based on the Araport11 annotation. Previously developed Python scripts were used to process the results and to generate final tables of raw counts, counts per million (CPMs), and fragment per kb per million (FPKMs) (58). Raw counts were subjected to DESeq2 (59) for identification of differentially expressed genes (DEGs) at adjusted p-value cutoff of 0.05. Genes without detectable expression in any sample were excluded from this analysis. Pairwise comparisons of *F*-m vs *f*-m, *f*-m vs *f*-K, *f*-m vs *f*-Q, and *f*-m vs *f*-R were performed, giving a fold change (FC) value. Only genes with log2FC >1 or <-1 were considered as DEGs. Heatmaps of DEGs were made by TBtool (60).

### Flavonol and camalexin quantification using LC-MS

Flavonol quantification through targeted metabolite analysis was performed according to the protocol described in previous studies (61, 62). Plant samples were grinded with liquid N_2_ and extracted with 80% methanol at room temperature. Supernatants were vacuum-dried in a SpeedVac at 65°C and pellets were redissolved in 80% aqueous methanol (1 μL mg^−1^ fresh weight). LC–MS analysis was done on an UPLC system (Exion LC Sciex) connected with a triple quadrupole system (QTRAP6500 +; ABSciex) using an electrospray ionization. Instrument settings and conditions were as described in Naik *et al.* (55). Analytical standards were obtained from Merck.

For camalexin quantification, 1 ml of 100% LC-MS grade methanol was added to the grinded plant samples. After centrifugation, the supernatant was taken out and filtered with 0.22 μm PVDF membrane filter Millex-GV (Merck Millipore) and samples were analyzed using the above LC-MS set-up. For quantification, a calibration curve was developed by measuring serial dilutions.

### Glucosinolate quantification using UHPLC

Glucosinolates were extracted from *A. thaliana* seedlings as desulpho-glucosinolates and analysed using high performance liquid chromatography (UHPLC) using established protocols (63, 64). Briefly, glucosinolates were extracted using 10 mg of freeze-dried sample in 70% methanol, containing 50 μM sinalbin as internal quantification standard. Desulfation of glucosinolates was performed overnight using purified sulfatase (25 mg ml^−1^; Sigma-Aldrich) on a DEAE Sephadex-A25 column. Desulfo-glucosinolates were eluted with 1 ml of HPLC-grade water and 20 μl of eluent was analysed using a Shimadzu CLASS-VP V6.14 HPLC machine and Luna C_18_ reverse-phased column (250 mm × 4.6 mm; 0.5 mm i.d.) through a gradient of water (solvent A; Merck-Millipore) and acetonitrile (solvent B; Merck-Millipore) gradient (0–1 min, 1.5% B; 1–6 min,1.5–5% B; 6–8 min, 5–7% B; 8–18 min, 7– 21% B; 18–23 min, 21–29% B; 23–23.1 min, 29–100% B; 23.1–24 min 100% B and 24.1–30 min 1.5% B; flow rate of 1.0 ml min^−1^). Glucosinolate components were detected at 229 nm and peaks were identified with reference to the standards. Data given are the averages ± standard deviation, of three independent measurements.

### Statistical tests

One-way ANOVA with Tukey’s post hoc test was applied for comparison of more than two datasets (*P*< 0.05). Each data set consists of three replicates.

## Results

### Feeding of flavonol derivatives to *A. thaliana* seedlings change metabolite levels

As a first step towards our venture to investigate the role of flavonols as regulators of gene expression and to analyse the potential role of flavonol derivatives as signaling molecules, we studied their uptake, and modification *in planta*. Flavonoids biosynthesis pathway in Arabidopsis and the nomenclature for the experiments are represented in Fig. 1A-B. We used the feeding of flavonol derivatives to seedlings of a flavonol-deficient *A. thaliana fls1-3* mutant to bypass the genetic absence of the FLS function. After feeding for 24h, washing and harvesting, LC-MS analysis was carried out to quantify the flavonol derivatives in the treated seedling samples (Fig. 1C). Kaempferol, quercetin, and rutin treatment caused significant increase in the respective derivative in both, Col-0 *(F)* and the *fls1-3* mutant *(f)*, indicating that flavonol aglycons as well as the flavonol glycoside (rutin) are taken up by the plants.

**Fig. 1.**
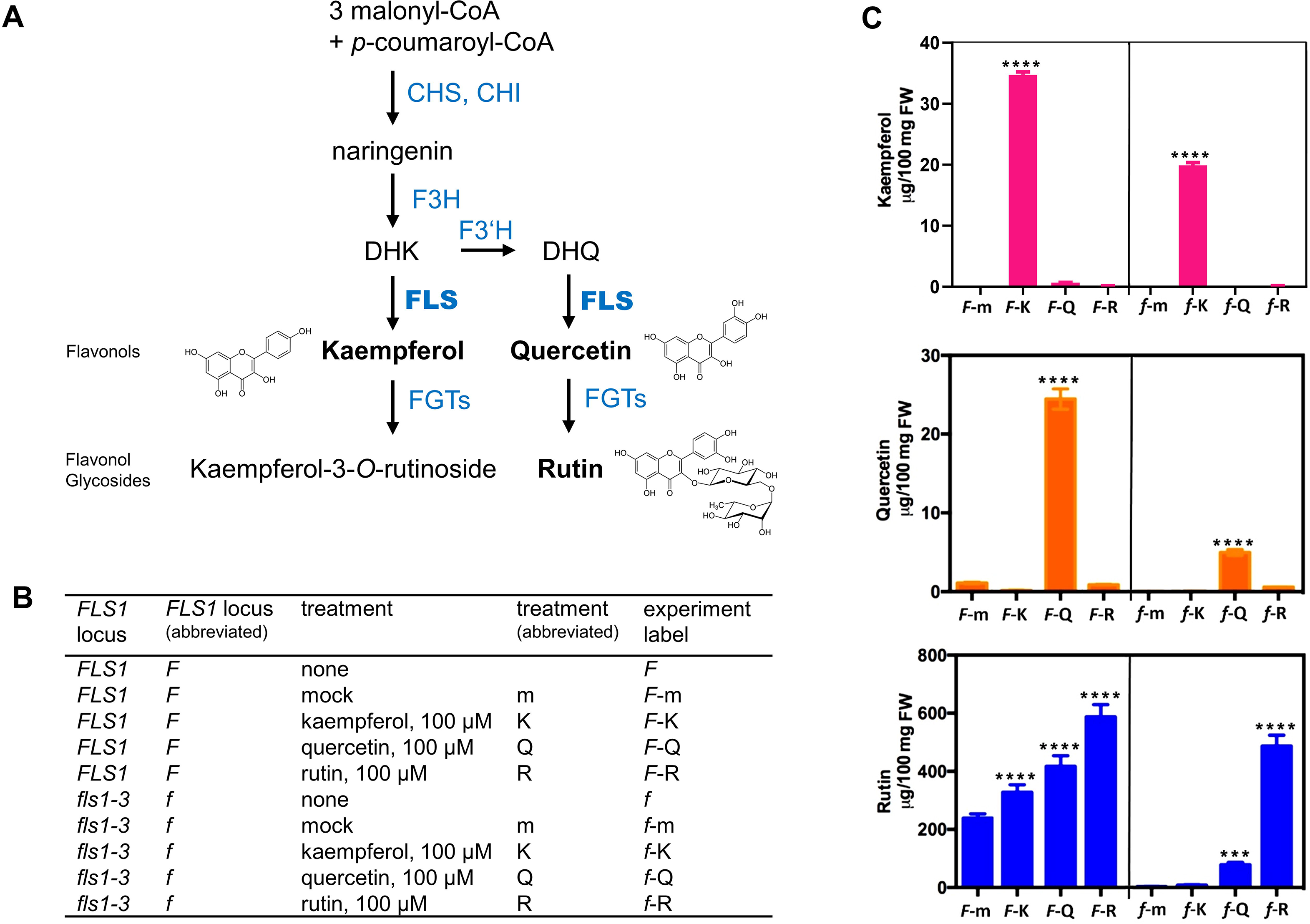
The supplement of exogenous flavonols affect flavonol metabolites levels in *Arabidopsis thaliana* seedlings. **(A)** Simplified schematic of the *A. thaliana* flavonoid pathway showing selected enzymatic steps (blue font) and metabolites (black font). Bold text identifies flavonol derivatives and the flavonol synthase enzyme. Enzyme names are abbreviated: chalcone synthase (CHS), chalcone isomerase (CHI), flavanone 3-hydroxylase (F3H), flavonoid 3′-hydroxylase (F3′H), flavonol synthase (FLS), flavonol glucosyl transferase (FGT). **(B)** Table represents the nomenclature for the experiments presented in this study. **(C)** Flavonol metabolite levels in 7 days old Col-0 *(F)* and *fls1-3 (f) A. thaliana* seedlings, fed for 24 h with 100 μM kaempferol (K), 100 μM quercetin (Q), 100 μM rutin (R) or mock (m). Values are presented as mean ± SD of the given metabolite content in μg/100 mg seedlings fresh weight from 2-3 replicates. Asterisks represent the level of significant change, where p-values <0.0001 (****) and between 0.0001 to 0.001 (***) denote extremely significant changes, between 0.001 to 0.01 (**) very significant and between 0.01 to 0.05 (*) significant changes.

### Chemical complementation of *fls1-3* mutant alters the expression of flavonol biosynthesis related genes in an antagonistic way

Next, in order to study a probable transcriptional regulation of flavonol biosynthesis due to the modulated content of flavonol derivatives, we analysed and compared in an targeted approach the transcript levels of regulatory and structural flavonol biosynthesis genes in *F*-m, *f*-m, *f*-K, *f*-Q, and *f*-R seedlings. The expression analysis of the R2R3-MYB-type flavonol regulators encoding genes revealed significant up-regulation of *MYB12* and *MYB111*, but not of *MYB11* in the *f*-m in comparison to *F*-m (Fig. 2). *MYB12*, the predominant flavonol regulator in seedlings roots, showed significant decrease in transcript abundance after treatment with all the three flavonol derivatives i.e., *f*-K, *f*-Q, and *f*-R. *MYB111*, expressed mainly in shoots, also showed significant down-regulation in *f*-K and *f*-R (Fig. 2). Structural genes of flavonol aglycon biosynthesis, like *CHS*, *CHI*, and *FLS* showed insignificant changes in their expression in the *f-*m mutant, however, a significant reduction was observed in *CHS* and *CHI* expression in both, *f*-K and *f*-R, as compared with *f*-m (Fig. 2). Also, the transcript levels of selected flavonol UDP-glycosyltransferases encoding genes were also found to be significantly increased in different combinations. For example, the expression of the flavonol-7-O-rhamnosyltransferase *UGT89C1* was upregulated in *f*-Q as compared with *f*-m. The expression of the flavonol-3-O-glycoside-7-O-glucosyltransferase *UGT73C6* was found to be significantly enhanced in all treatments (*f*-K, *f*-Q and *f*-R). However, the flavonol-3-O-rhamnosyltransferase *UGT78D1* displayed some up-regulation only in *f*-K and some downregulation in *f*-R (Fig. 2). Thus, flavonol deficiency and supplementation modulate the expression of genes involved in the flavonol biosynthesis and --glycosylation, suggesting the involvement of a feedback type of regulatory mechanism to fine-tune flavonol biosynthesis.

**Fig. 2.**
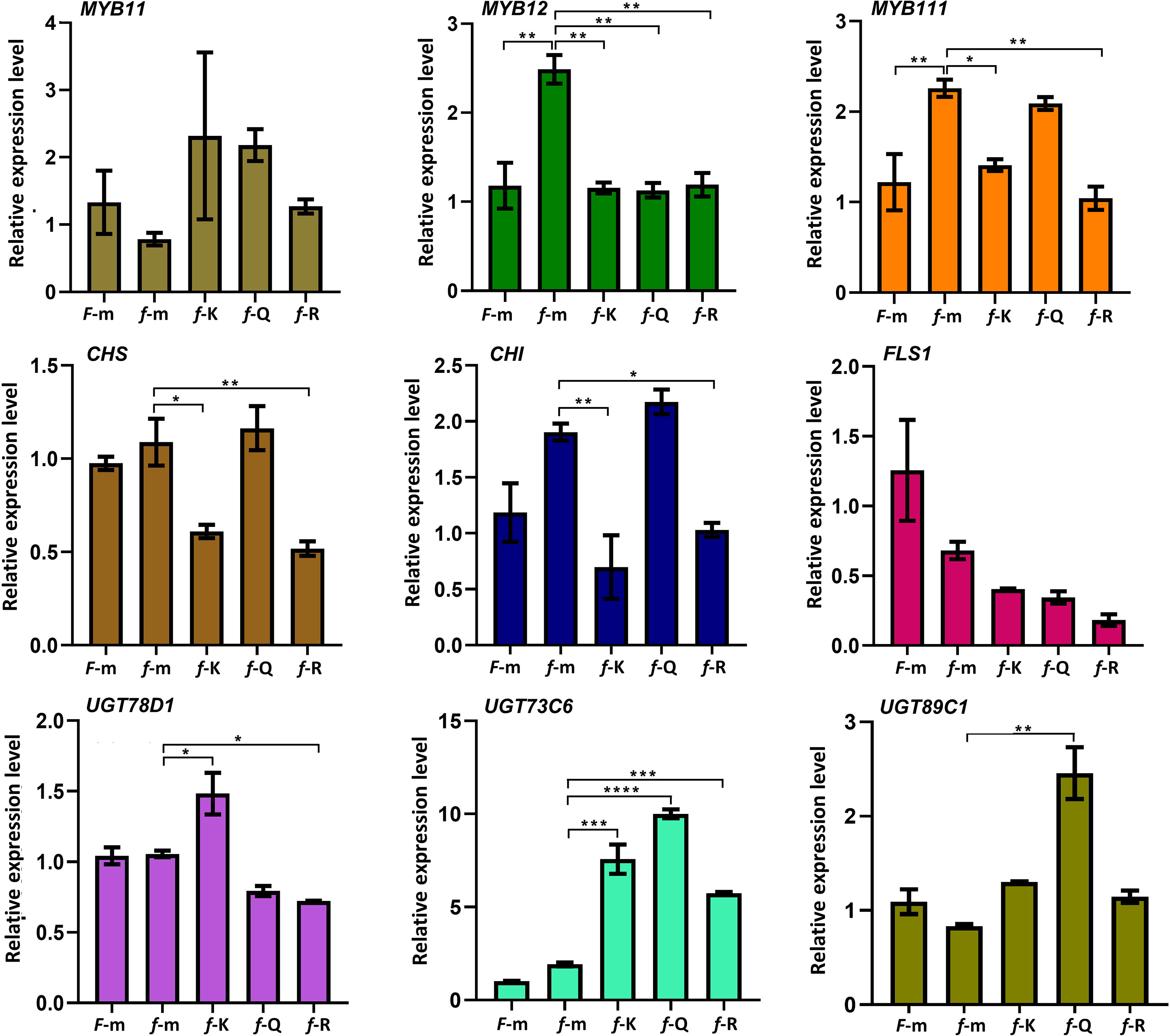
Expression analysis of flavonol biosynthesis related genes in seedling-feeding experiments. Relative expression of flavonol regulatory genes (*MYB11*, *MYB12*, *MYB111*) and structural biosynthesis genes (*CHS*, *CHI*, *FLS*, *UGT78D1* encoding a flavonol-3-O-rhamnosyltransferase, *UGT73C6* encoding a flavonol-3-O-glycoside-7-O-glucosyltransferase, *UGT89C1* encoding a flavonol-7-O-rhamnosyltransferase) in *F-*m, *f-*m, *f-*K, *f-*Q and *f-*R experiments, determined by RT-qPCR. Data is given as mean ± SD of triplicates. Significance levels are indicated with asterisks, where p-values <0.0001 (****) and between 0.0001 to 0.001 (***) denote extremely significant changes, between 0.001 to 0.01 (**) very significant and between 0.01 to 0.05 (*) significant changes.

### Transcriptome analysis reveals modulation of diverse pathways in the *fls1-3* mutant

To investigate the effect of flavonol deficiency over the global transcriptome, high-throughput RNA–Seq of the *F*-m, and *f*-m samples was carried out. In total, 474 genes were differentially expressed in *f*-m with log2FC >1 or <-1, as compared to *F*-m (Fig. 3A). Based on their annotations, the DEGs were found to be involved in diverse biological processes, including phytohormone biosynthesis and signalling, stress-response and specialized metabolism (*SI Appendix* Table S2, Fig. 3B). These modulations, therefore, can be attributed to the deficiency of flavonols. Thus, flavonol derivatives appear to affect various biological processes either directly or indirectly. Among these processes, we focused our study on the genes of the metabolic pathways, particularly those which are involved in specialized metabolism. To this end, as observed in the section 2 by RT-qPCR analysis (Fig. 2), the genes involved in the flavonol biosynthesis were modulated in the *f*-m as compared with *F*-m. In addition, the genes involved in suberin biosynthesis and deposition were found to be upregulated in *f-*m as compared to *F*-m seedlings (Fig. 3C). One of the notable observations was that the genes involved in camalexin and glucosinolate (GSL) biosynthesis process were differentially regulated in *f*-m (Fig. 3C). Although genes involved in the biosynthesis of both aliphatic and indolic GSLs were found to be differentially regulated, most of the DEGs were concerned with aliphatic GSL biosynthesis (Fig. 3C). Based on these observations, we assumed that flavonol deficiency might positively regulates genes involved in the suberin (*CYP86B1, GPAT5, ABCG6, FAR5, NAC58* and *CYP86A1*) and camalexin biosynthesis (*MYB122, NAC042, PAD3, CYP71A12* and *CYP79B2*), and negatively regulates the expression of genes involved in the aliphatic GSL (*IMD1, BAT5, MAM1, IPMI2, CYP79F2, IPMI1, CYP83A1, CYP79F1* and *BCAT4*) biosynthesis process. We also found a positive correlation between the expressions of glucosinolate biosynthesis genes and glucosinolate catabolism genes. Myrosinases (thioglucoside glucohydrolase), which are known to degrade GSL into isothiocyanates during biotic stress, also showed similar expression trends with the GSL biosynthesis genes. The expression of *BGLU37* (*TGG2*), *BGLU27*, and *BGLU28* decreased significantly in *F*-m vs *f*-m, while opposite trend was observed in *f*-m vs *f*-K, *f*-m vs *f*-Q, and *f*-m vs *f*-R (Fig. 3B).

**Fig. 3.**
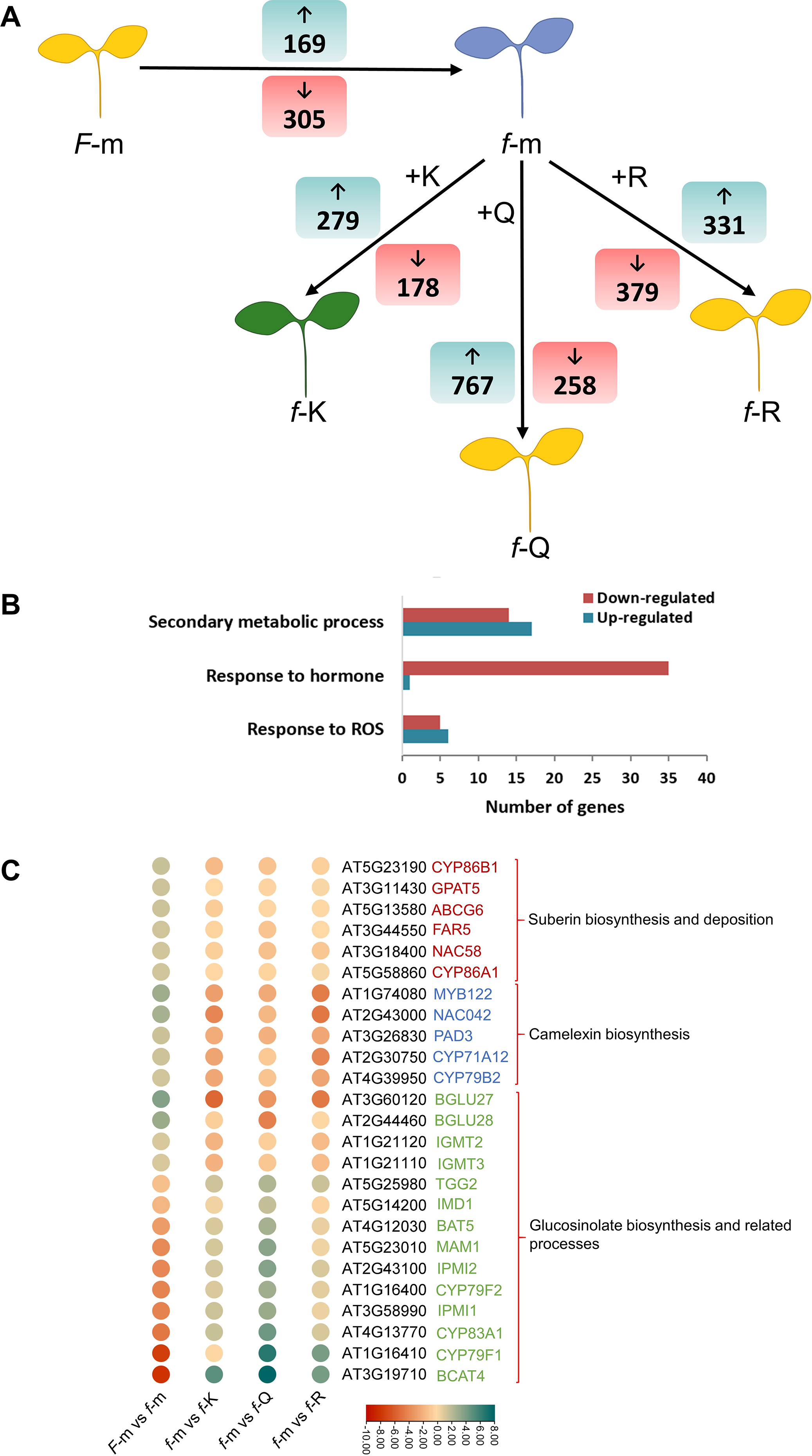
Transcriptome modulation by flavonol-feeding of seedlings. **(A)** Pictorial representation of flavonols seedling-feeding experiment showing the number of differentially expressed genes (DEGs) genes with log2FC ≥1 and log2FC ≤-1 at adjusted p-value >0.05 in *F*-m vs. *f*-m; *f*-m vs. *f*-K; *f*-m vs. *f*-Q; and *f*-m vs. *f*-R. Up- and down-regulated genes are shown with ↑ and ↓ arrows, respectively. Different colours of seedling are according to DPBA staining (Appelhagen *et al.,* 2014), indicating the accumulation of quercetin glycosides (yellow), kaempferol glycosides (green) or no flavonols (blue). **(B)** Bar graph showing the number of up- and downregulated DEGs involved in secondary metabolic processes, the response to hormones and to ROS in *F*-m vs. *f*-m. **(C)** DEGs from *fls1-3* mutant feeding-experiments involved in suberin biosynthesis and deposition, camalexin biosynthesis and glucosinolate biosynthesis and related processes. Color bar scale display low (dark-red) to high (aqua-blue) gene expression.

Further, we sought whether complementation with flavonols can modulate the gene expression in the *f-*m mutant. To this end, our transcriptome analysis revealed that a total of 457 (279 log2FC >1 and 178 log2FC<-1), 1025 (767 log2FC >1 and 258 log2FC<-1) and 710 (331 log2FC >1 and 379 log2FC<-1) genes were differentially expressed in *f*-K, *f*-Q, and *f*-R, respectively (*SI Appendix* Table S2; Fig. 3A). The expression of suberin biosynthesis and deposition-related genes were nearly not affected by the flavonoid feeding of the *fls1-3* mutant (*SI Appendix* Table S2). Genes involved in camalexin and aliphatic GSL biosynthesis but very well, so we studied these in detail. In general, chemical complementation reversed the expression pattern of these genes, when compared with their expression pattern observed in *F*-m vs *f-*m (Fig. 3B). For examples, several genes involved in the camalexin and aliphatic GSL biosynthesis were observed to be upregulated following the treatment with K, Q and R, which were otherwise downregulated in *f*-m as compared to *F*-m.- We conclude that flavonol treatment affects the expression of key genes involved in the camalexin and aliphatic GSL biosynthesis. Another notable observation was that treatment with different flavonol derivatives modulated the gene expression concerning with these pathways differentially. Only quercetin feeding appeared to modulate the expression of GLS biosynthesis-related genes in *fls1-3*. Maybe this hints to a quercetin specific functionality.

### Flavonol deficiency might positively regulate camalexin biosynthesis

To support the effect of different flavonol derivatives on camalexin biosynthesis at transcriptional level, we performed RT-qPCR expression analysis of key genes involved in the camalexin biosynthesis in *F*-m, *f-*m, *f-*K, *f-*Q, and *f-*R. RT-qPCR expression patterns in *F*-m vs *f-*m were in accordance to the expression trends observed in RNA-Seq data, as the transcript levels of the analysed genes were significantly upregulated in *f-*m (Fig. 4A-B). For example, the gene encoding a major rate-limiting enzyme of camalexin synthesis, *PHYTOALEXIN DEFICIENT3 (PAD3)*, was significantly upregulated in *f-*m. After flavonol feeding, particularly with quercetin and rutin, *PAD3* expression decreased substantially. A similar modulation of expression was seen for *CYTOCHROME P450 FAMILY 79 SUBFAMILY B2 (CYP79B2)* (Fig. 4B). The expression of *CYP79B3* showed significant reduction after kaempferol and quercetin feeding, while *GAMMA-GLUTAMYL PEPTIDASE1 (GGP1)* expression was reduced after quercetin and rutin treatment. The expression pattern of MYB122, the transcriptional regulator of camalexin biosynthesis, aligned well with the expression patterns of the camalexin biosynthesis genes (Fig. 4B). The expression of the NAC transcription factor *JUNGBRUNNEN1*/*NAC042* follows the expression patterns of its target genes (Fig. 4B). Further, in order to ascertain whether these transcriptional changes, as observed in the *f-*m and flavonol-treated *fls1-3 A. thaliana* seedlings are reflected at the metabolite level, we quantified camalexin through UHPLC-MS. The metabolite analysis showed manifold increase in the accumulation of camalexin in *f-*m as compared to *F*-m. Kaempferol, quercetin, and rutin treatment, on the other hand, substantially reduced the camalexin content (Fig. 4C). The presented results strongly suggest that flavonols are negative regulators of camalexin biosynthesis in *A. thaliana*.

**Fig. 4.**
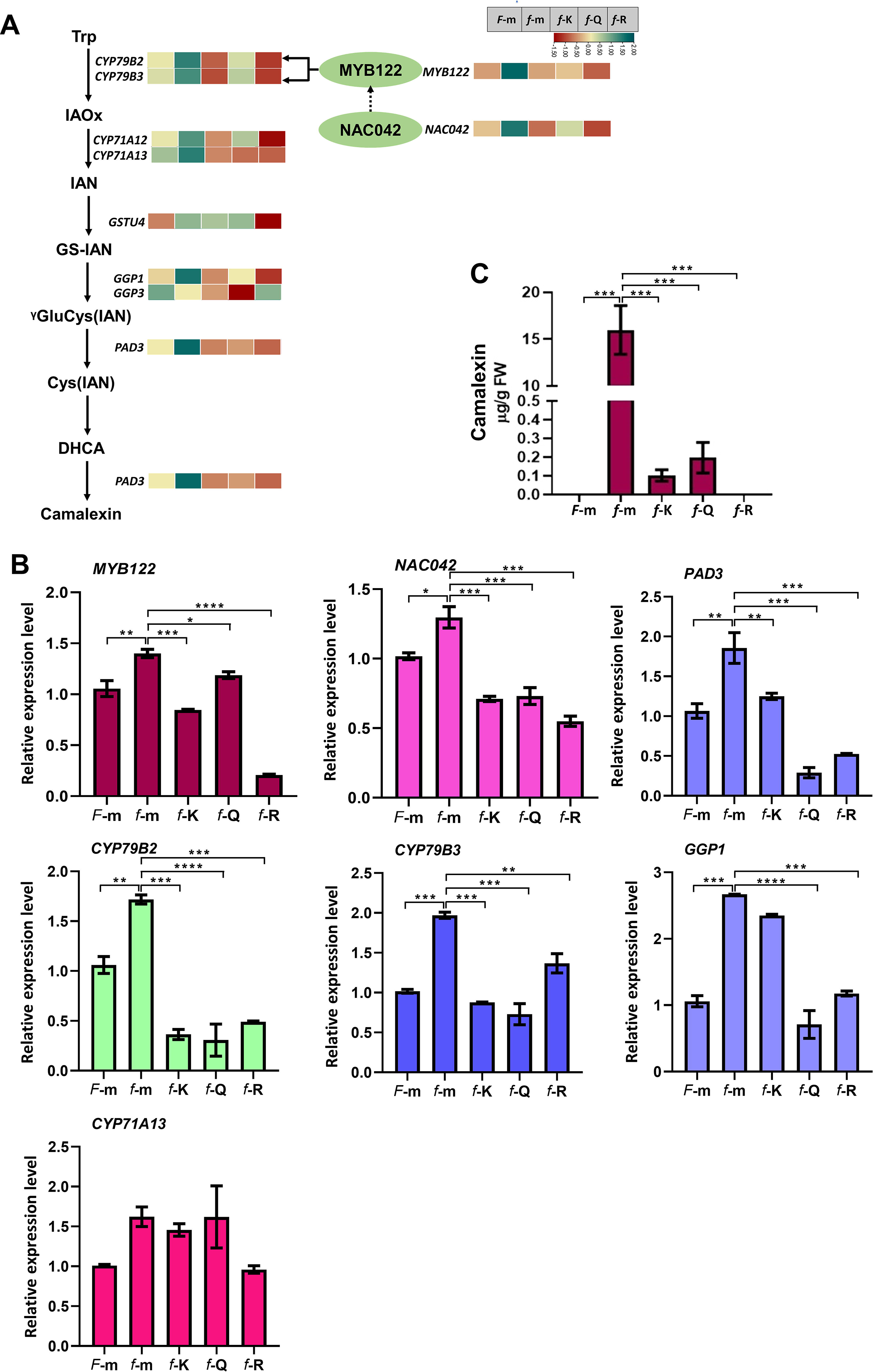
Flavonol-dependent expression profiling of camalexin pathway related genes and metabolite quantification. **(A)** A schematic pathway of camalexin biosynthesis in *A. thaliana* showing the expression of camalexin pathway related genes. Abbreviations: tryptophan (TRP), indole 3 acetaldoxime (IAOx), IAN (indole 3 acetonitrile), GS (glutathione), γGluCys (γ-glutamylcysteine), DHCA (dihydrocamalexic acid). The transcription factor MYB122 regulates the expression of *CYP79B2* and *CYP79B3*; and NAC042 indirectly regulates the expression of *MYB122*. Heatmaps give RNA-Seq data using normalized FPKM values of genes in *F*-m, *f*-m, *f*-K, *f*-Q, and *f*-R experiments (from left to right). Color bar scale displaying dark-red to aqua-blue colour shades corresponds with low to high gene expression. **(B)** RT-qPCR based relative expression level (referred to *F*-m) of various selected camalexin biosynthesis and regulatory genes. **(C)** Quantification of camalexin. Values represented the mean ± SD of three replicates. Asterisks denote the level of significance, where p-values <0.0001 (****) and between 0.0001 to 0.001 (***) denote extremely significant changes, between 0.001 to 0.01 (**) very significant and between 0.01 to 0.05 (*) significant changes.

### Glucosinolate biosynthesis is modulated by flavonol deficiency and flavonol complementation

RNA-Seq data of our flavonol-feeding experiments showed modulation in the transcript level of many biosynthesis genes involved in aliphatic GSL metabolism (Fig. 3C, Fig. 5A). Genes involved in aliphatic GSL biosynthesis were significantly modulated in the *fls1-3* mutant and after quercetin feeding. *CYTOCHROME P450* genes like *CYP79F1*, *CYP79F2*, and *CYP83A1* were down regulated in *f-*m, while flavonol treatment, particular with quercetin, could significantly increase their expression level. The encoded proteins of other biosynthesis genes like *METHYLTHIOALKYLMALATESYNTHASE1* (*MAM1*), *ISOPROPYLMALATEDEHYDROGENASE1* (*IMD1*), *2-ISOPROPYLMALATESYNTHASE2* (*IMS2*), *ISOPROPYLMALATEISOMERASE* (*IPMI*), *BRANCHED*-*CHAINAMINOTRANSFERASE4* (*BCAT4*) are responsible for GSL side chain elongation. Their expression was found to be decreased in *f*-m, while supplementation with different flavonols, again particularly with quercetin, induced their expression levels (Fig. 5A). Apart from the structural genes identified through differential transcriptome analysis, we also studied the expression pattern of two genes, encoding known transcriptional regulators of aliphatic GSL biosynthesis, *MYB28* and *MYB29*. The expression of *MYB28* and *MYB29* was comparatively reduced in *f-*m. However, *MYB29* transcript level significantly increased after quercetin treatment. Interestingly, the expression of *MYB28* was found to be decreased further after treatment with different flavonol derivatives. To validate this, we performed RT-qPCR analysis of *MAM1*, *CYP79F1*, *CYP79F2*, and *CYP83A1,* which supported the RNA-Seq data. Their expression was lower in *f-*m as compared to *F*-m (Fig. 5B). Flavonol treated *f*-K, *f*-Q and *f*-R had modulated transcript abundance of these genes than *f-*m. Thus, it can be assumed that flavonol metabolites positively regulate aliphatic GSL accumulation.

**Fig. 5.**
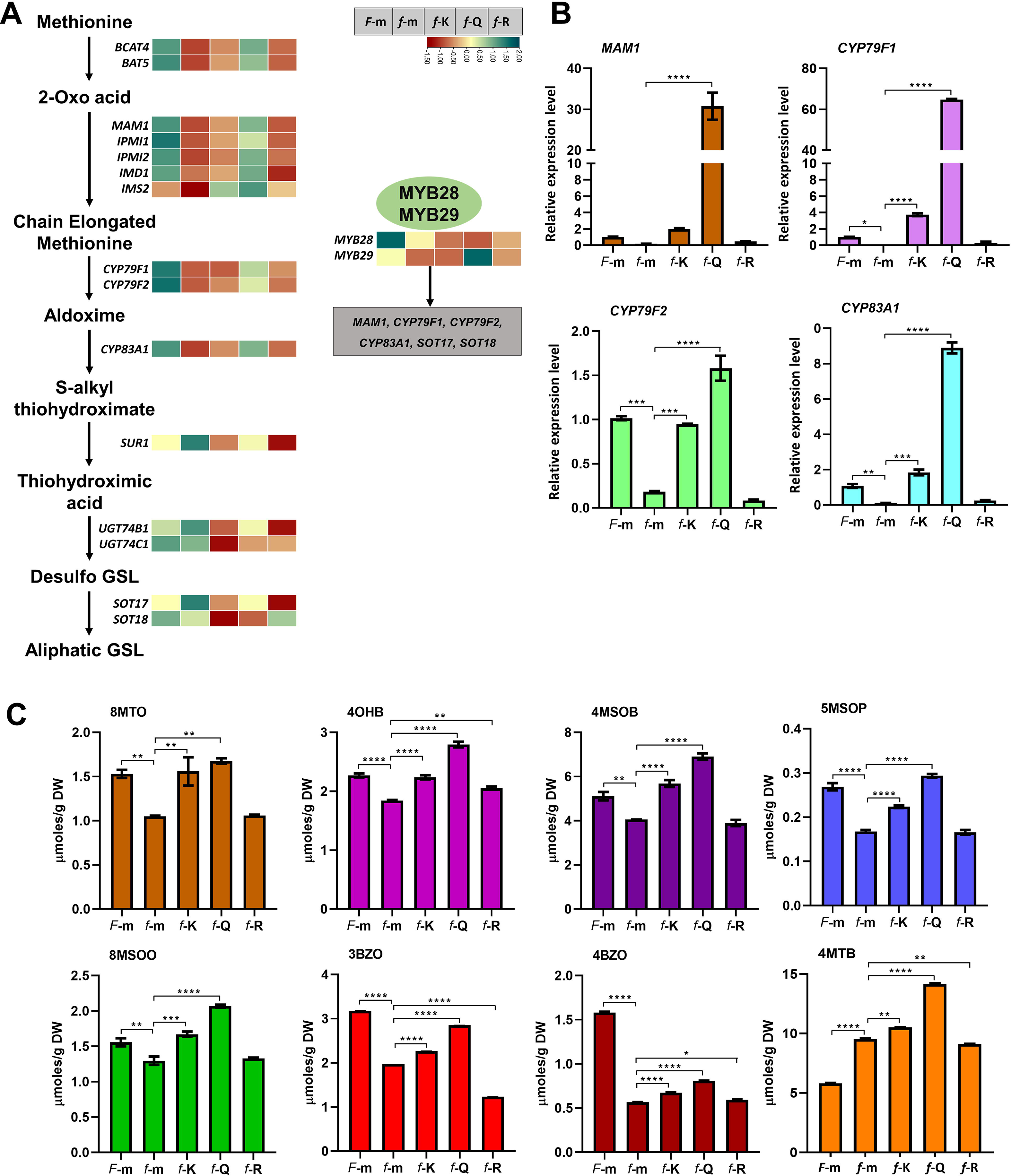
Flavonols promote aliphatic glucosinolates (GSL) synthesis. **(A)** A schematic pathway of aliphatic GSL biosynthesis in *A. thaliana* along with the RNA-Seq-derived heatmaps showing normalized FPKM values as measure for gene expression level in *F*-m, *f*-m, *f*-K, *f*-Q, and *f*-R experiments (from left to right). Dark-red to aqua-blue colour shades in color bar scale correspond with low to high gene expression. **(B)** RT-qPCR based relative expression level of various aliphatic GSL biosynthesis genes (referred to *F*-m). **(C)** Content of major aliphatic GSLs in *F*-m, *f*-m, *f*-K, *f*-Q, and *f*-R (8MTO, 8-methylthiooctyl; 4OHB, 4-hydroxybutyl; 4MSOB, 4-methylsulfinylbutyl; 5MSOP, 5-4- methylsulfinylpentyl; 8MSOO, 8- methylsulfinyloctyl; 3BZO, 3-benzoyloxypropyl; 4BZO, 4-methyloxybutyl; 4MTB, 4-methylthiobutyl). Values are given as mean ± SD of triplicates and the level of significant differences are marked with asterisk, where p-values <0.0001 (****) and between 0.0001 to 0.001 (***) denote extremely significant changes, between 0.001 to 0.01 (**) very significant and between 0.01 to 0.05 (*) significant changes.

To study the alteration in the GSL metabolome, we performed HPLC analysis to quantify major aliphatic GSL. Comparative metabolite analysis showed decreased accumulation of various methylated, benzoylated, hydroxylated, sulfinylated GSL in *f*-m (Fig. 5C; *SI Appendix* Fig. S1). Significantly reduced aliphatic GSLs in *f*-m include 4-hydroxybutyl (4OHB), 4-methylsulfinylbutyl (4MSOB), 7-methylsulfinylheptyl (7MSOH), 8-methylsulfinyloctyl (8MSOO), 3-benzoyloxypropyl (3BZO), 4-methyloxybutyl (4BZO), 7-methylthioheptyl (7MTH), and 8-methylthiooctyl (8MTO). On the other hand, chemical complementation in the *f* mutant enhanced the accumulation of different GSLs differentially. Quercetin treatment enhanced most of the aliphatic GSLs followed in the strength of the effect by kaempferol and rutin, respectively. Overall, our experiments suggest that different flavonol metabolites have divergent effects on the accumulation of variant aliphatic GSLs by modulating the expression of different biosynthetic genes. Quercetin seems to be the major modulator of aliphatic GSL synthesis in Arabidopsis.

## Discussion

Flavonols have been implicated in a variety of functions in plant biology ranging from mediating ecological interactions to serving as modulators of plant development (4, 5, 8, 10). Increasing reports on their involvement in plant growth and development as well as in stress response, along with their ubiquitous occurrence across land plants suggest that these small organic molecules functions beyond the classical definition of specialized metabolites. It is still an open question why there is such a variety of different flavonol derivatives and what specific functions they might have. In order to approach the answer to these questions, we thought that a detailed picture of transcriptome changes following the treatment of plants with flavonols has the potential to provide useful insights into diverse pathways regulated by these molecules. Therefore, we used an *A. thaliana* flavonol deficient mutant and identified several genes, which are differentially regulated by flavonol deficiency as well as by complementation with different flavonol derivatives.

In order to investigate, modulation of transcriptome following flavonol treatments, it is imperative to study flavonol uptake in Arabidopsis from growth media in a particular experimental set up and growth conditions. To this end, the phytochemical analysis suggested that different flavonols are taken up into seedlings from outside the media. This observation is in close accordance with the earlier study, which demonstrated active uptake of different flavonoids including dihydroflavonols, flavonol aglycones and flavonol glycoside (20, 27). The active transport of flavonoids, for example their efflux and vacuolar sequestration, has been reported to be mediated by ABC family of transporters (20, 21, 24, 65). The transcriptome resource developed in the present study should be useful in mining of candidate transporter genes in the future. Further, our complementation study revealed that kaempferol and quercetin treatment led to the accumulation of rutin along with the respective flavonol aglycones in Arabidopsis seedlings, suggesting that that flavonol aglycones tend to get modified into glycosylated forms. UDP-glycosyltransferases, such as *UGT89C1* and *UGT73C6*, catalyze rhamnosylation and glucosylation of quercetin and kaempferol at the 7-OH position, respectively (66, 67), while UGT78D1 catalyze rhamnosylation of flavonol at 3-OH position. In our study, we found that the expression of certain UDP-glycosyltransferases was significantly induced following quercetin and kaempferol treatment. These glycosyltransferase genes might be involved in the glycosylation of flavonols into respective flavonol derivatives. Also, the expression of structural and regulatory genes involved in the biosynthesis of flavonol aglycone backbone was found to be activated in the flavonol deficient mutant. Similar to these results, the expression of genes involved in the biosynthesis of flavonols and anthocyanins in the Arabidopsis *tt5/chi* seedlings, deficient in naringenin, was increased (25). Thus, owing to the defects in the specific steps in the flavonoid pathway, an unknown mechanism of feedback regulation might be operative to fine-tune the metabolism. Such feedback regulation of biosynthetic genes by pathway intermediates was first recognized, when the effect of flavonoid biosynthesis mutants on the accumulation of flavonoid mRNAs (68) and proteins (69) was investigated.

Our transcriptome analysis of *f-*m mutant and *F*-m seedlings revealed several differentially regulated genes, which were not just confined to the flavonoid biosynthesis, but involved in other pathways, such as hormone signaling, stress-response, suberin biosynthesis, and specialized metabolism. Previously, microarray-based global gene expression analysis of sucrose-induced, hydro-cultured *A. thaliana tt5/chi* mutant seedlings compared with *tt5/chi* seedlings complemented with naringenin (the product of CHI), suggested that genes involved in different pathways like stress-response and jasmonate signaling are modulated by flavonoids (25). Since naringenin is an intermediate leading to the biosynthesis of different sub-classes of flavonoids like flavonols, anthocyanins and PAs, the study by Pourcel et. al. (25) could not provide specific information about flavonol responsive genes, also because the experiment was conducted under anthocyanin-inducing conditions. In other studies, analysing the effect of treatments with individual flavonoid molecules such as kaempferol, quercetin, scutellarin, hyperoside were reported to alter the expression level of various genes as well as activity of several enzymes, resulting in the modulation of morpho-physiological parameters. For example, quercetin treatment in hydroponic cultured rice seedlings inhibited root growth and ROS level, and resulted in huge changes in the transcriptome (26). In Sorghum, flavonoid aglycones and glycosides modulated the expression of genes like *SHORT-ROOT, HD ZIP-III* and other root differentiation-related genes (27). The treatment of hyperoside (quercetin 3-O-galactoside) induced the expression of *CDPK6*, *MYB30*, *PAL2*, *CHS4*, *CHI3*, *FLS1*, and *UF3GaT1* in *Abelmoschus esculentus* (okra) (70). It is worth to mention that certain flavonoid derivatives display a restricted spatial distribution in the plant. However, flavonols like kaempferol, quercetin, and rutin are ubiquitously present in many different plant species (71, 72), and therefore appear to play important role in plants. Therefore, we decided to use kaempferol quercetin, and rutin in our experimental complementation approach to unravel the genes and pathways, which are specifically affected by each of these three flavonols.

One plausible mechanism to explain modifications of gene expression by flavonols stems from the antioxidant property of flavonols. Besides their damaging effect, ROS also function as signaling molecules, which modulate gene expression to allow for mitigation of oxidative stress (73–75). In view of this, we speculate that modulation of gene expression by flavonol deficiency, and complementation with flavonol metabolites might be mediated by alteration in the levels of ROS. Apart from their antioxidant activity, kaempferol, quercetin, rutin, and other flavonoids have been reported to either activate or inhibit diverse families of kinase enzymes, which are the active components of different developmental signaling pathways in animal (76–79). In plants, such mechanism has not been explored comprehensively; however, few recent studies have shown their direct effect on protein activity. For example, flavonol negatively modulate PINOID (PID) kinase, which is known to positively regulate PIN protein localization (32). PIN proteins play important roles in the regulation of auxin gradients, and therefore, flavonols can interfere with auxin transport (80–82). Thus, modulation of auxin transport might also contribute to the changes in the gene expression following perturbation of flavonol biosynthesis. In fact, flavonol molecules have long been known to inhibit polar auxin transport, and as compared to kaempferol, quercetin has been reported to be a more potent inhibitor of polar auxin transport (83). Studies have also suggested possible involvement of flavonols as inhibitors of auxin oxidation by ROS, and thereby these molecules can influence the level of active auxin in the cell (84). In addition, there are some indications that quercetin can influence ABA signaling (85). Taken together, it can be assumed that different flavonol derivatives may modify the auxin transport, auxin oxidation, ABA signaling and ROS levels differentially, which might result in their differential impact on the gene expression. Based on comparative transcriptome analysis of Col-0 and the *tt4-11/chs* mutant, in a very recent study (6), circadian rhythm was found to be influenced by flavonoids. Quercetin has been particularly tested for its role on the expression of the major clock gene *CIRCADIAN CLOCK ASSOCIATED1 (CCA1)*, where its supplementation enhances the promoter activity of *CCA1* in both Col-0 and the *tt4-11/chs* mutant (6). Therefore, flavonols, particularly quercetin, might be involved in the regulation of circadian rhythm, which in turn can affect gene expression.

One of the interesting observations of the present study is the inverse relationship between flavonol and camalexin biosynthesis (Fig. 4C). Camalexin, which is nearly undetactable in the flavonol containing wildtype, drastically accumulate in the flavonol-deficient *fls1-3* seedling and is reduced by flavonol treatment. Many genes involved in camalexin synthesis, particularly *MYB122*, *CYP71A13*, *CYP79B2*, *CYP79B3*, and *PAD3* were down-regulated upon flavonol treatment. Conversely, these genes were reported to be highly expressed in the flavonol deficient mutant. MYB122 acts as transcriptional activator of early steps in camalexin biosynthesis involving *CYP79B2*, and *CYP79B3* genes (40, 41). NAC042/JUB1, showed slight up-regulation in flavonol deficiency and down-regulation after flavonol supplementation. NAC042 has been shown to be involved in the induction of camalexin biosynthesis, possibly through modifying the expression of *CYP71A12*, *CYP71A13*, and *CYP71B15*/*PAD3* genes (86). Further, H_2_O_2_, one of the ROS molecules was reported to induce the expression of *NAC042* (87). Another study also revealed that H_2_O_2_ production induces the biosynthesis of camalexin in Arabidopsis (88). Camalexin biosynthesis is also regulated by two other classes of transcription factors ERF1, ERF72 and WRKY33, which are activated by MPK3/MPK6 (51, 52, 89) and ROS have been implicated in the activation of MPK3 and MPK6 in *A. thaiana* (30). In view of this, we speculate that modulation of camalexin biosynthesis by flavonols might be mediated by ROS, MPK3/MPK6 and NAC042 (Fig. 6). Detailed studies are required to dissect the molecular mechanism of negative cross-talk of camalexin and flavonol biosynthesis. Notably, NAC042 represses gibberellins (GAs) and brassinosteroids (BRs) biosynthesis by directly repressing the expression of the biosynthesis genes *GA3ox1* and *DWARF4*, respectively (90). Further, GA and BR signaling negatively regulate flavonol synthesis (11, 91). Therefore, the role of GA and BR signaling in the modulation of camalexin biosynthesis via flavonols needs further exploration.

**Fig. 6.**
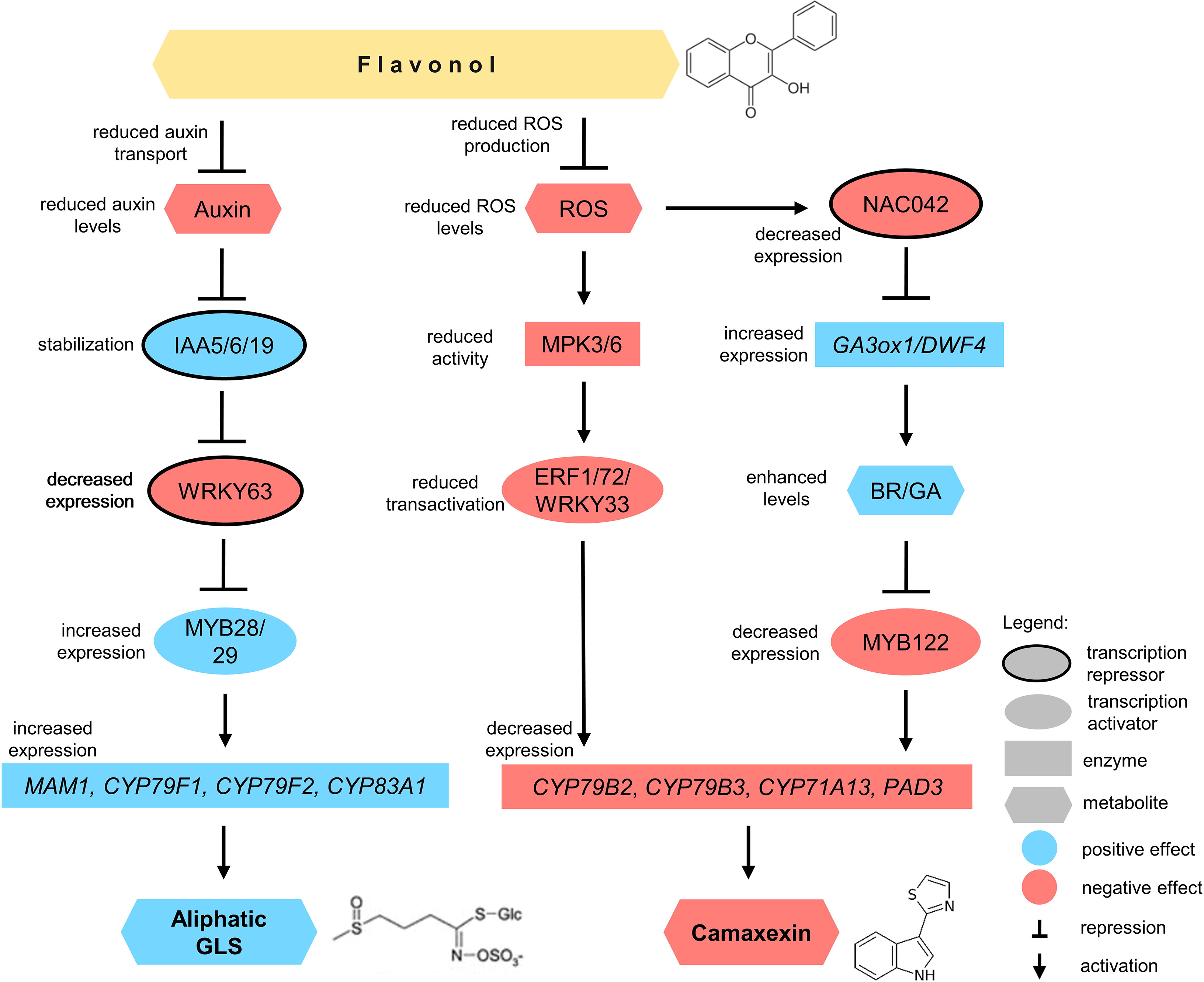
Mechanistic model of flavonol-induced modulation of camalexin and aliphatic glucosinolate biosynthesis in *Arabidopsis thaliana*. Flavonol reduced auxin transport and accumulation which results in stabilization of IAA5, IAA6, and IAA19 repressor. It leads to decreased expression of *WRKY63*, and subsequently increased expression of its target *MYB28* and *MYB29.* It promotes aliphatic glucosinolate synthesis via up-regulation of expression of its biosynthesis genes like *MAM1*, *CYP79F1*, *CYP79F2*, and *CYP83A1*. Flavonol also inhibit ROS accumulation and subsequently decreased MPK3/6 activity. Decreased MPK activity leads to reduced transactivation of ERF1, ERF72, and WRKY33, and ultimately decreased camalexin accumulation. Decreased ROS level after flavonol treatment also causes decreased *NAC042* expression, which may lead to increased expression of *GA3ox1* and *DWF4*, and increased gibberellin and brassinosteroids level. Increased GA and BR levels decreased the expression of *MYB122*, *CYP79B2*, *CYP79B3*, *CYP71A13* and *PAD3* and decreased camalexin content. Positive and negative regulators are depicted in blue and red boxes, respectively.

Another notable observation of the present study is the modulation of GSL biosynthesis by flavonols. Based on the gene expression and phytochemical analysis, a positive relationship between aliphatic GSL and flavonol biosynthesis is seen. In our study, quercetin was found to be the major inducer of numerous aliphatic GSL biosynthesis genes followed by kaempferol (Fig. 5B). The mechanism may involve auxin and the downstream signaling apparently (Fig. 6). Inhibition of auxin transport results in reduced accumulation of auxin, which stabilizes the transcriptional repressor proteins INDOLE-3-ACETIC ACID INDUCIBLE5 (IAA5), IAA6 and IAA19 (92). The stabilized IAA proteins in turn repress the expression of *WRKY63*, the negative regulator of *MYB28* and *MYB29* (92). Repression of WRKY63 leads to higher expression of *MYB28* and *MYB29*, subsequently higher expression of its biosynthesis genes target like *MAM1*, *CYP79F1*, *CYP79F2*, and *CYP83A1* etc. Auxin negatively regulates aliphatic GSL biosynthesis via this module. Flavonol probably promotes aliphatic GSL synthesis via inhibition of auxin accumulation. However, the mechanism by which flavonols impact GSL biosynthesis remains to be studied. Again, the ROS scavenging activity of flavonol presumably plays an important role in this type of metabolic cross-talk. This possibility is further supported by an earlier study which demonstrated that ROS induces indolic GSL biosynthesis (88). Since flavonols have been reported to inhibit auxin transport and might prevent auxin oxidation (93, 94), the role of auxin in regulating such metabolic changes cannot be ruled out.

As evident by the number of genes modulated by flavonols, it can be concluded that different flavonols differentially affect the gene expression. For example, out of the three flavonols tested in the present study, quercetin has been reported to be the most potent one. Thus, different flavonol molecules display differential bioactivity in the plant system. Phytohormones like auxin and gibberellins exists in structurally different forms, which display varying bioactivities (51, 95–98). These structural variants and modified forms assure homeostasis of these molecules and allow for the induction of signaling cascades in proper intensity as per the requirement. The instances of occurrence of structurally diverse flavonol and their further modifications in plants could be therefore viewed analogous to the structural diversity within a particular phytohorome. In view of this, presence of structurally diverse flavonols helps in fine-tuning the regulations offered by them.

Taken together, the present study reveals a detailed picture of genome-wide modulation in the gene expression by different types of flavonol metabolites. Our study suggested that flavonols, might function as signaling molecules and modulate the biosynthesis of other classes of specialized metabolites. The modulation of diverse pathways by flavonols as demonstrated in the present study, further strengthens the view that flavonols are not merely specialized metabolites, but these molecules offer phytohormone-like activity to the plant system. Our study should motivate further investigations to establish detailed mechanism of regulation of plant growth, development and metabolism by flavonols.

## Acknowledgements

This work was supported by the core grant of National Institute of Plant Genome Research (NIPGR) and Department of Biotechnology grant (BT/PR/38402/GET/119/308/2020) to AP. JN and RR acknowledge Council of Scientific and Industrial Research, Government of India, for Senior Research Fellowships. ST acknowledges Department of Biotechnology, for Research Associate (RA) Fellowship. The authors are thankful to DBT-eLibrary Consortium (DeLCON) for providing access to e-resources. We are grateful to Christoph Ringli, University of Zurich for providing the *A. thaliana fls1-3* mutant. We acknowledge the Metabolome facility (BT/INF/22/SP28268/2018) at NIPGR for phytochemical analysis. Many thanks to the German network for bioinformatics infrastructure (de.NBI, grant 031A533A) and the Bioinformatics Resource Facility (BRF) at the Center for Biotechnology (CeBiTec) at Bielefeld University for providing an environment to perform the computational analyses.

## Conflict of interest statement

Authors declare no conflict of interest.

## Data availability

All data supporting the findings of this study are available within the paper and within the supplementary data published online. RNA-Seq data sets are available from the Sequence Read Archive under BioProject accession number PRJNA880681.

## Author contributions

AP conceived the idea and designed the research. JN, ST, RR, and PK performed the experiments. BP did the computational analyses and identified differentially expressed genes (DEGs) in pairwise comparisons. PM analysed the RNA-Seq data. NCB helped in execution of glucosinolates quantification. JN, ST and PM wrote the initial draft, RS, and AP revised the manuscript. All the authors have read and approved the manuscript for submission.

## Supporting information

**Fig. S1. Quantification of aliphatic glucosinolates.** Content of some aliphatic GSLs including 3MSOP, 3-methylsulfinylpropyl; 7MTH, 7-methylthioheptyl; 5MTP, 5-methylthiopentylin; GNA, gluconapin; and 7MSOH, 7-methylsulfinylheptyl in *F*-m, *f*-m, *f*-K, *f*-Q, and *f*-R. Values are shown as mean ± SD of triplicates and the significant difference between *F*-m and *f*-m, and between *f*-m, and *f*-K, *f*-Q, and *f*-R, respectively, are marked with asterisks.

**Table S1.** List of primers used in the study.

**Table S2.** Differentially expressed genes in *F*-m vs *f*-m; *f*-m vs *f*-K, *f*-m vs *f*-Q and *f*-m vs *f*-R along with the baseMean, log2-foldchange, adjusted p-value and their annotation.

## Notes

### Competing Interest Statement

The authors have declared no competing interest.

