## Supplementary Information for "Flavonols contrary affect the interconnected glucosinolate and camalexin biosynthesis pathway in *Arabidopsis thaliana*"

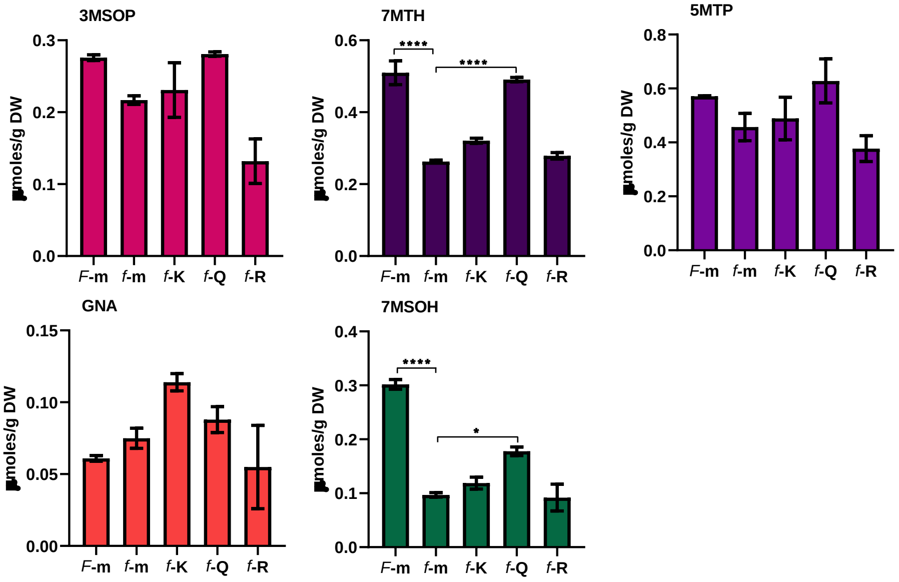


**Fig. S1. Quantification of aliphatic glucosinolates.** Content of some aliphatic GSLs including 3MSOP, 3-methylsulfinylpropyl; 7MTH, 7-methylthioheptyl; 5MTP, 5-methylthiopentylin; GNA, gluconapin; and 7MSOH, 7- methylsulfinylheptyl in *F*-m, *f*-m, *f*-K, *f*-Q, and *f*-R. Values are shown as mean ± SD of triplicates and the significant difference between *F*-m and *f*-m, and between *f*-m, and *f*-K, *f*-Q, and *f*-R, respectively, are marked with asterisks.

**Table S1. List of primers used in the study**

| Primer name | Primer sequence (5’-3’) | Specifications |
|---|---|---|
| MYB12_RTF | CCAAGTTAACGATGCGTCG | Forward primer for expression analysis of MYB12 |
| MYB12_RTR | GGCCTCATCATCACCGTC | Reverse primer for expression analysis of MYB12 |
| MYB111_RTF | TGGAAAGAGAAAGAGAGGGAAG | Forward primer for expression analysis of MYB111 |
| MYB111_RTR | CTCCCAATCAAGCAACTCC | Reverse primer for expression analysis of MYB111 |
| MYB11_RTF | CTCATGTGAGTCGAACAACG | Forward primer for expression analysis of MYB11 |
| MYB11_RTR | AAGGGTTTGACCTTCTCTCC | Reverse primer for expression analysis of MYB11 |
| CHS_RTF | GTGCTTCTTCTTTGGATGAG | Forward primer for expression analysis of CHS |
| CHS_RTR | GATGCGGAAGTAGTAGTCAG | Reverse primer for expression analysis of CHS |
| CHI_RTF | ATGTCTTCATCCAACGCC | Forward primer for expression analysis of CHI |
| CHI_RTR | CGCCGAGGAATAATGGATTG | Reverse primer for expression analysis of CHI |
| FLS_RTF | GGAGGTCGAAAGAGTCCAAGAC | Forward primer for expression analysis of FLS1 |
| FLS_RTR | AATCGCCGGCGTTGGAC | Reverse primer for expression analysis of FLS1 |
| UGT89C1_RTF | GAATGGTCGGAGGAGTTATGTTG | Forward primer for expression analysis of UGT89C1 |
| UGT89C1_RTR | GATGAGCGTCGTGTTAAAGAAATG | Reverse primer for expression analysis of UGT89C1 |
| UGT78D1_RTF | TAACAGATGCCTTCTTCTGG | Forward primer for expression analysis of UGT78D1 |

| | | |
|---|---|---|
| UGT78D1_RTR | CACTATCCCTTGCTCTCTTG | Reverse primer for expression analysis of UGT78D1 |
| UGT73C6_RTF | AGGTAGGAGTAGACAAAGCAGA | Forward primer for expression analysis of UGT73C6 |
| UGT73C6_RTR | TCAATTATTGGACTGTGCTAGTTG | Reverse primer for expression analysis of UGT73C6 |
| NAC042_RTF | ATTGGAACCAACATATAGTTGG | Forward primer for expression analysis of NAC042 |
| NAC042_RTR | GATCTAAGTTCATCCCAACTTG | Reverse primer for expression analysis of NAC042 |
| MYB122 _RTF | ACCTCTTCGAATCTCCCCATC | Forward primer for expression analysis of AtMYB122 |
| MYB122 _RTR | AACTTCATTGATCGGCGTCAC | Reverse primer for expression analysis of AtMYB122 |
| PAD3 _RTF | CTTGCTCCCAAGACAGACAA | Forward primer for expression analysis of AtPAD3 |
| PAD3 _RTR | GATCACGACCCATCGCATAA | Reverse primer for expression analysis of AtPAD3 |
| CYP79B2 _RTF | GTAACTTCGGAGCATTCGT | Forward primer for expression analysis of AtCYP79B2 |
| CYP79B2 _RTR | TCGCCGGATATCACATCC | Reverse primer for expression analysis of AtCYP79B2 |
| CYP79B3 _RTF | AGTCACTTCCGAACACTCA | Forward primer for expression analysis of AtCYP79B3 |
| CYP79B3 _RTR | TCGCAGGTTACCATATTCC | Reverse primer for expression analysis of AtCYP79B3 |
| GGP1 _RTF | TCCTGACGAGAAAGATCTG | Forward primer for expression analysis of AtGGP1 |
| GGP1 _RTR | ACAATATCACAGAGCTTAAGG | Reverse primer for expression analysis of AtGGP1 |
| CYP71A13 _RTF | GGGTAGAGGCTGGACCAAAT | Forward primer for expression analysis of AtCYP71A13 |

| | | |
|---|---|---|
| CYP71A13 _RTR | ACAACCGAAGATGGAAATGC | Reverse primer for expression analysis of AtCYP71A13 |
| IGMT3 _RTF | GGCTTCTCCCATTGCCAGT | Forward primer for expression analysis of AtIGMT3 |
| IGMT3 _RTR | AAACTGAAATACAGATGAAATTAAC AAT | Reverse primer for expression analysis of AtIGMT3 |
| IGMT2 _RTF | CAAGTGTGAGAAGGTCTCCGTA | Forward primer for expression analysis of AtIGMT2 |
| IGMT2 _RTR | ACCACGTCTTTCAGTTGTGC | Reverse primer for expression analysis of AtIGMT2 |
| MAM1 _RTF | CGGCTGAAAGAGTTGGGATATG | Forward primer for expression analysis of AtMAM1 |
| MAM1 _RTR | TCGATGAAACCTGAGGAACTG | Reverse primer for expression analysis of AtMAM1 |
| CYP79F1 _RTF | CCATACCCTTTTCACATCCTACTAGT CT | Forward primer for expression analysis of AtCYP79F1 |
| CYP79F1 _RTR | GTAGATTGCCGAGGATGGGC | Reverse primer for expression analysis of AtCYP79F1 |
| CYP79F2 _RTF | ACTAGGATTTATCGTCTTCATCGCA | Forward primer for expression analysis of AtCYP79F2 |
| CYP79F2 _RTR | CTAGGACGAGTCATGATTAGTTCGG | Reverse primer for expression analysis of AtCYP79F2 |
| CYP83A1 _RTF | GACCGAACCCTGATGAGTTTAG | Forward primer for expression analysis of AtCYP83A1 |
| CYP83A1 _RTR | GAGGAGAAGGTTCGCATAAGG | Reverse primer for expression analysis of AtCYP83A1 |
| Actin_RTF | ATGACATGGAGAAGATCTGGCATCA | Forward primer for expression analysis of Actin (For reference gene) |
| Actin_RTR | AGCCTGGATGGCAACATACATAGC | Reverse primer for expression analysis of Actin (For reference gene) |

**Supplementary Table 2. Differentially expressed genes in *F* -m vs. *f***

## *F* -m vs. *f* -m

| ID | baseMean | l2fc | padj |
|---|---|---|---|
| AT3G47340 | 9285.980172 | -2.254493487 | 2.73E-94 |
| AT1G11260 | 10369.48553 | -1.848671535 | 4.60E-59 |
| AT1G12240 | 3156.667637 | -1.760992639 | 2.29E-53 |
| AT4G35770 | 16790.9552 | -1.888479142 | 3.69E-48 |
| AT1G03090 | 5469.160874 | -1.515181901 | 4.55E-48 |
| AT4G25580 | 620.836364 | -2.611685374 | 2.68E-46 |
| AT5G63160 | 12582.13856 | -1.681603891 | 1.26E-45 |
| AT5G56870 | 9792.019405 | -1.832295386 | 2.03E-44 |
| AT2G34430 | 12892.95454 | -1.842096813 | 6.78E-41 |
| AT1G10070 | 7540.487887 | -1.855500538 | 1.94E-39 |
| AT5G44120 | 140.9646295 | 5.583522607 | 4.39E-37 |
| AT4G36850 | 4825.241727 | -1.686587206 | 4.53E-37 |
| AT3G05727 | 1918.515705 | -2.024241294 | 2.56E-35 |
| AT5G41080 | 8437.903665 | -1.449228494 | 6.16E-33 |
| AT4G26260 | 1017.053533 | -1.885946245 | 5.92E-31 |
| AT1G75945 | 118.9778773 | -5.420504059 | 3.66E-30 |
| AT3G16770 | 7084.99079 | -1.338778727 | 8.90E-30 |
| AT1G02660 | 2653.022331 | -1.352039407 | 1.22E-29 |
| AT5G49360 | 16237.88271 | -1.189792178 | 1.40E-29 |
| AT1G21400 | 10400.04806 | -1.536660811 | 4.88E-28 |
| AT5G23010 | 125.4916216 | -3.98115436 | 1.33E-27 |
| AT1G08630 | 6174.993195 | -1.591914224 | 2.61E-27 |
| AT3G61198 | 505.4554944 | -2.543716537 | 2.64E-27 |
| AT5G27920 | 1211.932146 | -1.365472127 | 8.93E-26 |
| AT3G15450 | 37186.33405 | -1.035391312 | 1.27E-25 |
| AT4G32870 | 472.323423 | -2.184035971 | 8.08E-25 |
| AT1G80160 | 1005.118224 | -1.83283465 | 1.17E-23 |
| AT3G06850 | 2860.313356 | -1.315835065 | 6.52E-23 |
| AT5G09440 | 1684.688051 | -1.34414787 | 4.40E-22 |
| AT3G13450 | 2939.37657 | -1.334077754 | 7.71E-22 |
| AT5G07080 | 287.5154888 | -2.102343959 | 1.89E-21 |
| AT5G18170 | 4999.20994 | -1.373481244 | 1.89E-21 |
| AT1G77210 | 1252.261329 | -1.336783926 | 4.49E-21 |
| AT5G51970 | 5226.854254 | -1.007526371 | 4.49E-21 |
| AT3G45730 | 1611.481379 | -1.670083509 | 7.11E-21 |
| AT1G10340 | 659.5692551 | -1.827290145 | 1.68E-20 |
| AT4G37870 | 9409.776644 | -1.186725994 | 4.32E-20 |
| AT3G56360 | 5724.610814 | -1.171101322 | 1.13E-19 |
| AT4G34030 | 3542.52634 | -1.14726094 | 1.74E-19 |
| AT4G27450 | 855.4186275 | -1.394824714 | 2.21E-19 |
| AT3G19390 | 1274.253399 | -1.440545086 | 6.44E-19 |

| | | | |
|---|---|---|---|
| AT3G00800 | 110.5362637 | -3.149919768 | 1.32E-18 |
| AT4G15530 | 8603.284865 | -1.389486391 | 1.79E-18 |
| AT1G13300 | 1010.872109 | -1.467016974 | 1.87E-18 |
| AT1G19540 | 360.8829412 | -1.597321553 | 1.87E-18 |
| AT2G33830 | 17133.39241 | -1.028431963 | 3.09E-18 |
| AT5G02020 | 1428.749982 | -1.341189421 | 7.19E-18 |
| AT2G38400 | 2651.146983 | -1.09452815 | 1.29E-17 |
| AT1G66760 | 1485.558973 | -1.204499421 | 1.40E-17 |
| AT1G66180 | 8670.314243 | -1.651026941 | 2.36E-17 |
| AT2G17880 | 1119.711765 | -1.224744781 | 2.64E-17 |
| AT1G79360 | 177.2742119 | -2.446067936 | 3.42E-17 |
| AT1G15040 | 3318.432671 | -1.226813492 | 5.25E-17 |
| AT1G19530 | 2576.098675 | -1.135900397 | 6.80E-17 |
| AT3G61060 | 2249.740037 | -1.04832609 | 1.28E-16 |
| AT4G19160 | 3474.609771 | -1.266601221 | 1.71E-16 |
| AT2G39980 | 1269.388654 | -1.10029088 | 2.39E-16 |
| AT5G25980 | 999.2222396 | -1.433833207 | 5.12E-16 |
| AT5G64110 | 433.2336091 | -1.386208625 | 5.16E-16 |
| AT1G79700 | 911.6990949 | -1.206850714 | 8.68E-16 |
| AT1G22500 | 1084.981773 | -1.522949048 | 3.22E-15 |
| AT1G06570 | 2769.138467 | -1.027786164 | 1.01E-14 |
| AT5G21170 | 2238.82704 | -1.011556437 | 2.25E-14 |
| AT5G13930 | 1666.204253 | 1.171562595 | 6.26E-14 |
| AT5G24530 | 750.3158911 | -1.241231214 | 7.08E-14 |
| AT2G36120 | 2402.952181 | -1.180021979 | 1.70E-13 |
| AT3G29240 | 5629.743782 | -1.025525668 | 1.87E-13 |
| AT5G53970 | 1602.082778 | -1.233754635 | 5.70E-13 |
| AT5G21020 | 4589.213806 | -1.017596391 | 8.35E-13 |
| AT2G43620 | 2758.296668 | -1.137422576 | 8.90E-13 |
| AT4G28040 | 937.5723165 | -1.103960126 | 1.00E-12 |
| AT1G26770 | 2354.272011 | 1.03493127 | 1.36E-12 |
| AT3G20395 | 548.5311547 | -1.174836133 | 1.60E-12 |
| AT3G45300 | 4331.278013 | -1.105770659 | 3.00E-12 |
| AT1G55920 | 2241.065743 | 1.118035281 | 3.93E-12 |
| AT5G44585 | 1627.857304 | -1.455146615 | 5.62E-12 |
| AT2G36490 | 387.5845674 | -1.310866525 | 7.08E-12 |
| AT4G25630 | 1630.753228 | 1.009088779 | 7.40E-12 |
| AT5G39520 | 117.786018 | -2.839993405 | 1.55E-11 |
| AT4G39950 | 3252.370481 | 1.277300987 | 1.56E-11 |
| AT3G20440 | 818.009124 | 1.00049602 | 1.94E-11 |
| AT3G58990 | 44.56098314 | -4.262899425 | 2.18E-11 |
| AT2G32150 | 4469.32828 | -1.059530536 | 2.68E-11 |
| AT3G45290 | 384.8868956 | -1.429695494 | 3.98E-11 |
| AT5G09220 | 177.584451 | -1.654912015 | 5.13E-11 |
| AT2G44380 | 1075.706218 | -1.140370707 | 5.37E-11 |
| AT1G78370 | 293.4845076 | -1.770641456 | 5.57E-11 |
| AT3G21870 | 489.1501839 | -1.281536667 | 6.85E-11 |

| | | | |
|---|---|---|---|
| AT5G01740 | 214.7344499 | -1.496946277 | 7.71E-11 |
| AT5G09530 | 292.9240314 | 1.41888703 | 1.20E-10 |
| AT2G43100 | 40.80722154 | -4.016149337 | 1.31E-10 |
| AT5G05250 | 364.6660736 | -1.789062094 | 1.45E-10 |
| AT4G12500 | 1043.789775 | -1.540670868 | 1.89E-10 |
| AT5G25260 | 645.1165358 | -1.574693785 | 3.21E-10 |
| AT4G21990 | 3526.849994 | 1.04979852 | 4.32E-10 |
| AT1G15010 | 165.2780131 | -1.760039628 | 5.12E-10 |
| AT1G76690 | 688.6840664 | 1.027347569 | 5.12E-10 |
| AT4G28420 | 159.1470109 | 2.120883241 | 5.91E-10 |
| AT5G52300 | 129.9468685 | -2.109373338 | 6.38E-10 |
| AT3G48390 | 407.5309597 | -1.4408234 | 7.02E-10 |
| AT2G18193 | 193.5742217 | -1.505709729 | 7.59E-10 |
| AT4G33960 | 390.2220503 | -1.11457699 | 8.87E-10 |
| AT1G66700 | 136.3758829 | 2.330931987 | 9.04E-10 |
| AT5G56100 | 1890.904307 | -1.210877819 | 9.04E-10 |
| AT2G39450 | 1307.159958 | -1.172845699 | 1.22E-09 |
| AT5G14200 | 156.5070643 | -1.85137466 | 1.23E-09 |
| AT3G26830 | 278.5479119 | 1.459426462 | 1.80E-09 |
| AT1G02640 | 1779.28595 | -1.026554683 | 2.19E-09 |
| AT3G56400 | 243.2442869 | -2.017101772 | 2.37E-09 |
| AT5G13170 | 89.18200802 | -2.391358025 | 3.64E-09 |
| AT2G47460 | 104.8839484 | 1.947676967 | 3.75E-09 |
| AT1G17170 | 1664.866015 | 1.012213376 | 4.00E-09 |
| AT1G70830 | 196.0714695 | -1.48576581 | 4.66E-09 |
| AT2G03865 | 71.73562089 | -3.300701068 | 6.19E-09 |
| AT1G35670 | 1443.433885 | -1.030960467 | 6.27E-09 |
| AT1G17745 | 540.0917096 | 1.286965586 | 6.77E-09 |
| AT5G14180 | 1110.149728 | -1.053900387 | 6.85E-09 |
| AT2G40420 | 668.3611592 | -1.093205987 | 7.29E-09 |
| AT3G54600 | 182.7800353 | -1.566090279 | 9.39E-09 |
| AT3G28930 | 1153.499268 | 1.164823507 | 9.76E-09 |
| AT3G06355 | 773.5294397 | -1.048662202 | 1.02E-08 |
| AT3G60120 | 45.34997798 | 3.706105474 | 1.21E-08 |
| AT2G27380 | 33.77469754 | 4.14253304 | 1.28E-08 |
| AT3G02480 | 210.2184424 | -1.533043733 | 1.40E-08 |
| AT1G62580 | 170.2711774 | 1.489974471 | 1.88E-08 |
| AT5G55970 | 888.6499466 | -1.015472607 | 2.24E-08 |
| AT5G62920 | 140.3926992 | 1.718401897 | 2.24E-08 |
| AT5G49480 | 3029.616988 | 1.094969564 | 2.54E-08 |
| AT5G50915 | 370.0215077 | 1.161854974 | 2.82E-08 |
| AT3G19710 | 25.92603561 | -8.11429382 | 3.18E-08 |
| AT3G16150 | 736.9080698 | -1.308116418 | 3.77E-08 |
| AT1G29395 | 95.83953669 | -1.828534951 | 4.86E-08 |
| AT1G69252 | 888.9255716 | -1.071494963 | 6.67E-08 |
| AT2G44460 | 45.94490118 | 2.974930752 | 6.78E-08 |
| AT2G39800 | 904.2417055 | -1.08866912 | 8.11E-08 |

| | | | |
|---|---|---|---|
| AT1G52690 | 171.0494707 | -1.510355061 | 8.90E-08 |
| AT1G16400 | 33.1948127 | -4.213179638 | 9.28E-08 |
| AT2G19810 | 1093.330954 | -1.061833639 | 9.41E-08 |
| AT5G46730 | 161.2273104 | -1.565456899 | 1.47E-07 |
| AT5G52310 | 819.9883388 | -1.319665717 | 1.67E-07 |
| AT1G45201 | 370.0822807 | -1.266749252 | 1.81E-07 |
| AT5G15960 | 86.76703699 | -2.409702774 | 3.08E-07 |
| AT3G54640 | 5593.54081 | 1.004178811 | 3.44E-07 |
| AT2G41190 | 236.7180552 | -1.224192773 | 4.28E-07 |
| AT2G22470 | 886.1257545 | 1.173123824 | 5.13E-07 |
| AT1G74930 | 200.7956814 | -1.35948633 | 5.16E-07 |
| AT1G16410 | 17.8448747 | -7.575459649 | 7.38E-07 |
| AT3G02020 | 24.75410595 | -5.575073854 | 7.94E-07 |
| AT2G29490 | 991.9194166 | 1.014250929 | 8.47E-07 |
| AT3G23880 | 423.5497447 | -1.126933117 | 8.84E-07 |
| AT4G13770 | 120.4672304 | -4.837183302 | 1.01E-06 |
| AT1G33960 | 165.4240514 | -1.918136425 | 1.65E-06 |
| AT2G04050 | 313.7010117 | -1.051776225 | 1.85E-06 |
| AT2G44370 | 1364.18747 | -1.034763496 | 1.95E-06 |
| AT5G14580 | 360.7451131 | 1.060638341 | 2.02E-06 |
| AT3G15460 | 220.7092638 | 1.162570392 | 2.03E-06 |
| AT1G03400 | 339.1824698 | -1.085358173 | 2.23E-06 |
| AT1G19610 | 3004.435738 | -1.037460868 | 2.23E-06 |
| AT5G58860 | 161.7495932 | 1.286640262 | 2.23E-06 |
| AT5G11420 | 391.8228744 | -1.01483424 | 2.24E-06 |
| AT1G10585 | 238.4079658 | 1.453656422 | 2.24E-06 |
| AT1G21110 | 983.7879235 | 1.077556307 | 2.27E-06 |
| AT3G25250 | 121.4813129 | 1.679649113 | 2.56E-06 |
| AT2G40340 | 256.2009494 | 1.092794464 | 2.68E-06 |
| AT5G65140 | 343.6170343 | 1.102732372 | 2.68E-06 |
| AT4G28703 | 151.9023921 | -1.383955352 | 3.01E-06 |
| AT1G74710 | 340.3337534 | -1.185976144 | 3.23E-06 |
| AT5G57240 | 160.7575151 | -1.441844047 | 3.68E-06 |
| AT1G26240 | 14.60630816 | 7.351542076 | 3.89E-06 |
| AT3G52720 | 202.3709293 | -1.308486691 | 4.23E-06 |
| AT5G64100 | 2073.335824 | -1.044811128 | 4.26E-06 |
| AT5G05965 | 94.5683228 | -1.746642249 | 4.69E-06 |
| AT3G16670 | 1089.240272 | -1.045562161 | 5.51E-06 |
| AT1G13530 | 119.2357019 | -1.844635116 | 5.87E-06 |
| AT2G21820 | 19.99966773 | -4.830618178 | 5.95E-06 |
| AT1G74080 | 41.22395174 | 2.897978908 | 6.36E-06 |
| AT4G02845 | 334.6412915 | -1.096123355 | 6.91E-06 |
| AT5G14070 | 92.54091215 | -1.61509535 | 6.94E-06 |
| AT5G52900 | 323.5147387 | -1.004335278 | 7.29E-06 |
| AT1G06957 | 13.59204725 | -7.180506164 | 7.50E-06 |
| AT3G59550 | 107.1892875 | 1.452429393 | 8.06E-06 |
| AT1G68500 | 194.6493332 | -1.152631698 | 1.07E-05 |

| | | | |
|---|---|---|---|
| AT1G09420 | 307.391076 | -1.038136019 | 1.07E-05 |
| AT1G51860 | 295.5705095 | -1.196559925 | 1.09E-05 |
| AT3G61890 | 187.7388225 | -1.309323386 | 1.10E-05 |
| AT3G62950 | 383.4971077 | -1.37998346 | 1.20E-05 |
| AT5G13210 | 93.02140251 | -1.544212079 | 1.28E-05 |
| AT5G26920 | 1005.756849 | -1.018832994 | 1.34E-05 |
| AT1G52000 | 146.1962992 | -1.353983925 | 1.35E-05 |
| AT2G25460 | 252.3760688 | -1.246482604 | 1.39E-05 |
| AT2G19670 | 237.9264115 | 1.002992587 | 1.45E-05 |
| AT4G34250 | 165.999626 | -1.361564673 | 1.65E-05 |
| AT1G67980 | 236.5463142 | 1.426988257 | 1.83E-05 |
| AT2G41870 | 234.2328354 | -1.00040138 | 1.92E-05 |
| AT1G05340 | 647.605839 | -1.138301684 | 2.22E-05 |
| AT2G23540 | 156.1920486 | 1.16093408 | 2.23E-05 |
| AT5G40990 | 45.84555879 | 2.141822856 | 2.26E-05 |
| AT1G21250 | 163.3546272 | -1.824812539 | 2.54E-05 |
| AT3G10110 | 226.1765289 | 1.048907181 | 2.89E-05 |
| AT5G10180 | 584.1089859 | -1.00239362 | 3.25E-05 |
| AT2G48130 | 97.81081151 | 1.422964008 | 3.26E-05 |
| AT4G38080 | 225.6287415 | 1.147031265 | 3.72E-05 |
| AT1G48130 | 36.77217131 | 3.249862456 | 4.05E-05 |
| AT1G14950 | 13.60007148 | 7.248500046 | 4.31E-05 |
| AT3G05955 | 23.59845815 | -3.278759658 | 4.38E-05 |
| AT5G54740 | 18.6040777 | 4.833682551 | 4.54E-05 |
| AT5G25490 | 132.4923939 | -1.204308339 | 4.72E-05 |
| AT3G06780 | 134.9761306 | -1.306358761 | 5.16E-05 |
| AT3G22840 | 171.7460639 | 1.196095068 | 5.85E-05 |
| AT3G05400 | 124.1715795 | 1.283438411 | 6.43E-05 |
| AT5G42530 | 326.6009086 | -3.859146834 | 6.50E-05 |
| AT4G23280 | 160.6673801 | 1.312668469 | 6.66E-05 |
| AT4G28460 | 65.96867231 | 1.929302946 | 6.71E-05 |
| AT1G17180 | 74.74684533 | 1.832627459 | 6.85E-05 |
| AT5G66480 | 990.7058144 | -1.029116813 | 7.55E-05 |
| AT5G15120 | 117.3464187 | 1.390109493 | 7.63E-05 |
| AT2G02990 | 137.5426999 | 1.38572812 | 7.72E-05 |
| AT5G44260 | 446.6965647 | -1.016635676 | 7.82E-05 |
| AT3G09405 | 181.7116692 | 1.430409652 | 8.03E-05 |
| AT1G04800 | 101.3446833 | -1.576454452 | 8.45E-05 |
| AT4G08770 | 312.211655 | 1.090613701 | 8.66E-05 |
| AT5G26230 | 34.17231822 | -2.629969536 | 8.84E-05 |
| AT2G22510 | 68.73702913 | 1.737943507 | 9.18E-05 |
| AT2G42900 | 250.2574837 | -1.031378647 | 9.72E-05 |
| AT5G66400 | 1156.218436 | -1.965473673 | 9.72E-05 |
| AT1G78970 | 108.5439377 | -1.428780805 | 0.000110698 |
| AT5G02780 | 35.28316995 | 2.457933295 | 0.0001176 |
| AT1G74670 | 761.7933397 | -1.0355017 | 0.000118907 |
| AT4G27160 | 19.53421334 | 3.508874438 | 0.000118907 |

| | | | |
|---|---|---|---|
| AT1G56430 | 31.23971613 | -2.473824382 | 0.000136553 |
| AT4G23890 | 232.5442617 | -1.074892731 | 0.000171533 |
| AT5G10946 | 87.79034706 | -1.374339962 | 0.000181409 |
| AT1G07620 | 69.12685738 | 1.553218507 | 0.000183291 |
| AT1G17960 | 88.09134193 | -1.353905799 | 0.000184501 |
| AT5G22580 | 100.7400104 | -1.33926411 | 0.000188875 |
| AT3G49410 | 160.6918153 | 1.060774642 | 0.000207687 |
| AT5G03350 | 85.81857714 | -2.131454703 | 0.00021321 |
| AT4G18940 | 52.54533641 | -2.180299781 | 0.000225127 |
| AT2G14247 | 47.80509396 | -2.284220134 | 0.00022608 |
| AT4G27150 | 15.86618365 | 4.599284974 | 0.000230296 |
| AT2G47770 | 52.2356336 | -1.986296616 | 0.000237037 |
| AT3G47780 | 192.7802236 | 1.192535934 | 0.000243165 |
| AT2G05695 | 15.35949732 | -5.465993303 | 0.000252612 |
| AT5G17350 | 182.5268838 | -1.114519857 | 0.000253022 |
| AT4G20970 | 131.6972444 | 1.336385402 | 0.000258267 |
| AT1G67865 | 163.9571924 | -1.613925294 | 0.000271346 |
| AT5G65600 | 126.2602753 | 1.480775228 | 0.000279719 |
| AT1G55990 | 43.91487155 | 1.928457979 | 0.000288859 |
| AT1G19960 | 9.459374435 | -6.660850022 | 0.000288911 |
| AT2G32870 | 71.2271526 | -1.508724673 | 0.000290455 |
| AT1G16150 | 58.79247604 | -1.689014772 | 0.000311032 |
| AT1G53490 | 108.595796 | 1.182064312 | 0.000333617 |
| AT2G14160 | 28.0049101 | -2.288734729 | 0.000354219 |
| AT2G35380 | 68.6538731 | 1.449730088 | 0.000358583 |
| AT5G37690 | 38.97873415 | 2.08950052 | 0.000378806 |
| AT4G20390 | 102.6617078 | 1.299604051 | 0.000395451 |
| AT1G14880 | 9.176795089 | -6.616569984 | 0.000438708 |
| AT2G01422 | 13.83943705 | 4.392978317 | 0.000446209 |
| AT2G41280 | 34.75096897 | -2.130600135 | 0.000446209 |
| AT1G31580 | 746.6597489 | -2.591796025 | 0.000455175 |
| AT5G06530 | 162.0135895 | -1.082802869 | 0.000455175 |
| AT3G18400 | 81.35807462 | 1.300541251 | 0.000467875 |
| AT1G61440 | 52.20404327 | 1.757660216 | 0.000470002 |
| AT5G58390 | 44.07496016 | -1.895860291 | 0.000512657 |
| AT4G01450 | 143.6814154 | -1.032612703 | 0.000536856 |
| AT5G22200 | 18.79647033 | -3.333350554 | 0.000548517 |
| AT1G01110 | 51.99901902 | -1.865596441 | 0.000550322 |
| AT2G40370 | 35.59986565 | 2.096796855 | 0.00055352 |
| AT3G05770 | 20.30649201 | 2.81667691 | 0.000621803 |
| AT2G27690 | 40.15277049 | -1.827212123 | 0.000629351 |
| AT3G12710 | 213.0932787 | -1.02356642 | 0.000630298 |
| AT1G51140 | 123.8951106 | -1.15769471 | 0.000651267 |
| AT1G18710 | 18.6211025 | -3.021130151 | 0.000690815 |
| AT5G38940 | 195.1900749 | -1.058023177 | 0.000724908 |
| AT1G69920 | 223.7588912 | 1.23691745 | 0.000787551 |
| AT4G12030 | 22.15382778 | -3.057066924 | 0.000809833 |

| | | | |
|---|---|---|---|
| AT5G05960 | 26.02418223 | -2.506052596 | 0.000835592 |
| AT1G52450 | 141.966753 | 1.041075035 | 0.000868871 |
| AT3G50440 | 117.545767 | -1.054372585 | 0.000870927 |
| AT4G16000 | 88.82551044 | -1.424071214 | 0.000889763 |
| AT2G26010 | 24.38657333 | -3.207590202 | 0.000902685 |
| AT5G02890 | 107.556712 | -1.159006832 | 0.00091395 |
| AT1G20850 | 194.9288623 | -1.238874649 | 0.001106468 |
| AT1G64370 | 725.2544412 | -1.59308576 | 0.001133017 |
| AT1G51380 | 126.7170523 | 1.140358839 | 0.001141801 |
| AT1G67910 | 171.5235811 | -1.006776293 | 0.001176908 |
| AT4G37290 | 83.08115524 | 1.437601028 | 0.001300334 |
| AT1G60750 | 24.94527827 | 2.364209672 | 0.001351632 |
| AT5G38030 | 83.88638284 | 1.364486044 | 0.00145415 |
| AT1G60110 | 41.42411761 | -1.956328201 | 0.001475669 |
| AT5G38710 | 223.2603366 | -1.042250068 | 0.001520582 |
| AT3G16120 | 91.90182911 | -1.279439542 | 0.001525149 |
| AT2G26695 | 38.65298206 | -1.903555968 | 0.001601536 |
| AT1G15050 | 31.03115648 | -2.183632634 | 0.001626137 |
| AT5G09580 | 102.5276517 | 1.187587473 | 0.001673067 |
| AT4G27460 | 74.14371989 | -1.399041581 | 0.001800597 |
| AT3G49580 | 43.27733963 | 1.799825269 | 0.001887673 |
| AT4G17030 | 113.9569864 | -1.299292348 | 0.001889644 |
| AT3G14580 | 79.60533401 | 1.202807455 | 0.001893785 |
| AT5G19600 | 107.0446256 | -1.304029607 | 0.001941108 |
| AT1G68850 | 56.24744749 | 1.655800836 | 0.002029069 |
| AT3G06520 | 136.4872955 | 1.050414257 | 0.002029069 |
| AT3G28580 | 31.43986752 | 2.179753727 | 0.002029069 |
| AT3G22740 | 20.76444754 | -2.532348001 | 0.002083029 |
| AT5G06930 | 72.8112093 | -1.233427303 | 0.002252717 |
| AT1G16850 | 36.10126047 | -2.023918345 | 0.002296926 |
| AT1G07400 | 96.31188628 | 1.08750941 | 0.002528273 |
| AT5G03615 | 260.944444 | -1.006687887 | 0.002550736 |
| AT5G60730 | 61.08870835 | 1.441904675 | 0.002594628 |
| AT3G21370 | 22.82433353 | 2.235332633 | 0.002736423 |
| AT3G50800 | 51.44984351 | -1.581176163 | 0.002780313 |
| AT4G33610 | 52.66965881 | 1.56752583 | 0.002792991 |
| AT5G57550 | 34.26393216 | -2.404863172 | 0.002829479 |
| AT1G34670 | 52.03270417 | 1.434859006 | 0.003242648 |
| AT3G25882 | 44.55177503 | -1.884799235 | 0.003291823 |
| AT4G23150 | 19.77961617 | 2.573651388 | 0.003361116 |
| AT1G76960 | 23.42306631 | -2.560158144 | 0.003428405 |
| AT5G48900 | 155.0767549 | -1.164244944 | 0.003729435 |
| AT1G19510 | 9.832571159 | -4.807513408 | 0.003760375 |
| AT4G21650 | 10.4096972 | -3.843562864 | 0.004056356 |
| AT5G59320 | 363.9909778 | -1.966765735 | 0.00449027 |
| AT5G12030 | 61.13027135 | 1.616528593 | 0.004591163 |
| AT2G17036 | 48.97392454 | -1.673304806 | 0.004599706 |

| | | | |
|---|---|---|---|
| AT2G05895 | 18.97435712 | 2.58751938 | 0.004617858 |
| AT5G13580 | 47.68485666 | 1.412411962 | 0.004737402 |
| AT4G22070 | 75.3090375 | 1.215119046 | 0.004742533 |
| AT4G30460 | 71.5774851 | -1.16539264 | 0.004879628 |
| AT3G55710 | 37.10143066 | -1.678382563 | 0.005100363 |
| AT1G30720 | 3804.489725 | -1.267866545 | 0.005204933 |
| AT3G53530 | 108.7783591 | -1.076055567 | 0.005313354 |
| AT5G35580 | 54.53718259 | 1.29031714 | 0.005372932 |
| AT2G45050 | 81.26979265 | -1.186093825 | 0.00553273 |
| AT1G64360 | 10.73125566 | -3.887826044 | 0.005831843 |
| AT1G13609 | 51.00540469 | -1.721522063 | 0.005881574 |
| AT5G04200 | 12.78332153 | -3.07035902 | 0.005881574 |
| AT1G27670 | 78.47117183 | -1.145889923 | 0.005952307 |
| AT5G37990 | 64.28481999 | -1.270303423 | 0.006230838 |
| AT5G42800 | 27.0873965 | -2.110445361 | 0.006345929 |
| AT4G04223 | 80.0743617 | 1.075907401 | 0.006452187 |
| AT5G62280 | 137.4746832 | -1.047735312 | 0.006470264 |
| AT1G64940 | 42.27402573 | 1.473663703 | 0.006901689 |
| AT2G40010 | 157.2228393 | 1.019035682 | 0.006903208 |
| AT5G15970 | 915.8469735 | -1.345098933 | 0.006958496 |
| AT2G43590 | 2297.176063 | 1.16261419 | 0.007029633 |
| AT1G74460 | 76.88089811 | 1.258542631 | 0.007117088 |
| AT1G67750 | 26.66943975 | -1.854227661 | 0.007209709 |
| AT3G21550 | 25.98686211 | -2.158602579 | 0.00731788 |
| AT2G43000 | 200.1472866 | 2.621646354 | 0.007557456 |
| AT3G16360 | 43.61433193 | -1.389042431 | 0.007773958 |
| AT1G22160 | 24.55896667 | -2.329985633 | 0.007826981 |
| AT2G34360 | 47.99185086 | 1.348127565 | 0.008054586 |
| AT1G60590 | 93.42814471 | -1.017251654 | 0.008473902 |
| AT2G16890 | 86.0156492 | 1.053643585 | 0.008522295 |
| AT4G27140 | 12.10396756 | 3.574555691 | 0.008613364 |
| AT3G11430 | 39.9538151 | 1.445529883 | 0.008755199 |
| AT1G66040 | 29.10415754 | -1.910106917 | 0.00940511 |
| AT1G15830 | 56.05934844 | -1.222762893 | 0.009602365 |
| AT3G01345 | 24.81954314 | 2.186568193 | 0.009630404 |
| AT3G48340 | 47.67135838 | -1.328426873 | 0.009641735 |
| AT2G48140 | 98.60967407 | 1.002650402 | 0.009711396 |
| AT3G01190 | 39.23630705 | 1.527967199 | 0.010225001 |
| AT5G51810 | 45.4643869 | -1.333481164 | 0.010225001 |
| AT3G54940 | 18.14607599 | 2.853276891 | 0.010225951 |
| AT2G42870 | 127.8262806 | -1.026232136 | 0.010471523 |
| AT2G25680 | 97.45396784 | -1.151959548 | 0.010541524 |
| AT1G72870 | 18.89824409 | -2.378029742 | 0.011350335 |
| AT5G60760 | 54.11366438 | 1.248424316 | 0.011687593 |
| AT4G30450 | 28.5085434 | -1.777754688 | 0.011824201 |
| AT5G35660 | 9.459271099 | 3.789448474 | 0.011824201 |
| AT1G49960 | 10.93338415 | 2.92381948 | 0.011903365 |

| | | | |
|---|---|---|---|
| AT2G23270 | 37.36060839 | 1.909511013 | 0.012175961 |
| AT2G32210 | 197.9917162 | -1.094213331 | 0.012175961 |
| AT5G57560 | 6318.472377 | -1.019201734 | 0.012657754 |
| AT3G21380 | 47.98275389 | 1.318843847 | 0.012657912 |
| AT2G30370 | 27.24876395 | -1.690869728 | 0.013850104 |
| AT4G04450 | 68.06692727 | 1.169982101 | 0.014013529 |
| AT5G24330 | 46.91638797 | 1.329915308 | 0.014267525 |
| AT1G26380 | 1631.811206 | 1.318433678 | 0.014421012 |
| AT1G17285 | 15.1110857 | 2.537020844 | 0.014623967 |
| AT3G10870 | 57.69675807 | -1.34539197 | 0.015488569 |
| AT3G59480 | 24.03370184 | -1.893818998 | 0.015844666 |
| AT5G18430 | 35.34392962 | -1.662139686 | 0.01602351 |
| AT2G32200 | 97.29688664 | -1.082563698 | 0.016073892 |
| AT1G18320 | 63.65062682 | 1.285283652 | 0.016208737 |
| AT5G43580 | 6079.747409 | -1.162056547 | 0.016330091 |
| AT1G74810 | 123.8344173 | 1.024309217 | 0.016519224 |
| AT1G11850 | 113.3075567 | -1.064937833 | 0.016748984 |
| AT5G07475 | 26.96988558 | 1.838380165 | 0.017208872 |
| AT4G28520 | 221.2998038 | 3.974108723 | 0.017494833 |
| AT2G00340 | 73.87314718 | 1.106212524 | 0.017876724 |
| AT3G04320 | 24.46203977 | -2.057073173 | 0.017882754 |
| AT5G21150 | 63.99425325 | 1.157014957 | 0.018508555 |
| AT3G50760 | 67.99492657 | 1.046647155 | 0.018856878 |
| AT4G22160 | 72.82508488 | -1.015618248 | 0.019174819 |
| AT5G23190 | 41.15479527 | 1.523372794 | 0.019240653 |
| AT3G15357 | 140.1648121 | 1.057060438 | 0.019422945 |
| AT1G03100 | 46.1924884 | 1.293403971 | 0.019451538 |
| AT2G22590 | 41.70312085 | 1.345567812 | 0.019920912 |
| AT1G10990 | 79.58582095 | 1.017457197 | 0.019957175 |
| AT1G09867 | 16.88852774 | 2.116644455 | 0.020293753 |
| AT5G23700 | 29.23294662 | -1.918898495 | 0.021987404 |
| AT3G18170 | 62.69605273 | -1.240463286 | 0.022083056 |
| AT4G23990 | 31.79378715 | -1.456837119 | 0.022956683 |
| AT5G15500 | 42.67397191 | -1.551633275 | 0.023020244 |
| AT3G16530 | 1312.390359 | 1.29561862 | 0.023942811 |
| AT5G16410 | 17.09882165 | -2.19925746 | 0.024371926 |
| AT4G33070 | 45.51333756 | 1.381022048 | 0.025449415 |
| AT1G63310 | 64.19380319 | -1.177232735 | 0.02601001 |
| AT5G10120 | 12.76111733 | 2.502280498 | 0.02601001 |
| AT3G14440 | 57.35934227 | -1.21092812 | 0.026142968 |
| AT1G09475 | 23.97445141 | 1.938970757 | 0.026493599 |
| AT4G30250 | 15.85222494 | 2.139580397 | 0.026794296 |
| AT2G46640 | 34.92965659 | -1.563404809 | 0.027293844 |
| AT5G43890 | 39.63558676 | -1.340824945 | 0.027376306 |
| AT5G17040 | 17.51642253 | 1.96134938 | 0.027984077 |
| AT5G42210 | 12.70210134 | -2.38763115 | 0.0280721 |
| AT3G09190 | 46.2186168 | -1.213430915 | 0.029023402 |

| | | | |
|---|---|---|---|
| AT1G02380 | 58.67339754 | 1.155621505 | 0.029410505 |
| AT3G55990 | 34.04013372 | -1.590518506 | 0.029766688 |
| AT1G13800 | 49.23292994 | 1.174328459 | 0.030766138 |
| AT5G43180 | 56.39374435 | 1.049317354 | 0.030766405 |
| AT1G51920 | 50.12584587 | 1.450929752 | 0.031104466 |
| AT5G24940 | 66.23830331 | -1.310939689 | 0.031118988 |
| AT1G06553 | 31.97518352 | 1.649870296 | 0.031160582 |
| AT4G15320 | 10.52372227 | -2.751890303 | 0.031358243 |
| AT1G47980 | 18.81002001 | 1.948407447 | 0.031446293 |
| AT4G26120 | 95.09025078 | 1.02970276 | 0.031684975 |
| AT3G49142 | 74.12253993 | 1.008725583 | 0.032657102 |
| AT3G59710 | 15.06558106 | -2.180836731 | 0.032657102 |
| AT3G00190 | 14.94150862 | -2.664425007 | 0.032726427 |
| AT1G62540 | 21.8988353 | -1.986704168 | 0.032949884 |
| AT3G00390 | 20.28776475 | -1.84570633 | 0.032974706 |
| AT4G26990 | 81.87964521 | 1.00065796 | 0.033597936 |
| AT4G19870 | 84.88350103 | -1.117881934 | 0.034097001 |
| AT5G61250 | 71.46387522 | -1.051719042 | 0.034744255 |
| AT2G19970 | 62.9123452 | -1.245385537 | 0.035271246 |
| AT5G04150 | 31.20948527 | -1.456563781 | 0.037315986 |
| AT4G14120 | 44.15255443 | 1.390470091 | 0.037838933 |
| AT2G26290 | 41.78762882 | 1.268230454 | 0.03820443 |
| AT1G30730 | 1571.338238 | -1.085402926 | 0.038807014 |
| AT4G13690 | 41.42462493 | 1.199974962 | 0.038927939 |
| AT4G23160 | 19.2515737 | 1.788404275 | 0.039522679 |
| AT4G26960 | 65.88793252 | 1.018720297 | 0.039537757 |
| AT2G42540 | 444.2050206 | -2.140796526 | 0.039670551 |
| AT1G01680 | 62.11305694 | -5.154573148 | 0.040133122 |
| AT3G50400 | 32.65106271 | 1.354556366 | 0.040315623 |
| AT1G74490 | 22.49086305 | 1.62614996 | 0.041234647 |
| AT4G21020 | 11.81683184 | 2.654281953 | 0.0419495 |
| AT3G45130 | 140.7885203 | -1.021104433 | 0.042077314 |
| AT5G44582 | 72.14289803 | -1.102982105 | 0.042082267 |
| AT4G28720 | 32.14758191 | 1.35193486 | 0.043446277 |
| AT1G02335 | 16.47537781 | -2.236277775 | 0.044056523 |
| AT3G00850 | 45.35230903 | -1.415561113 | 0.044253381 |
| AT3G09480 | 76.87671827 | 1.023825177 | 0.044434171 |
| AT5G64870 | 35.78381681 | -1.400243852 | 0.044494783 |
| AT3G51540 | 56.67137716 | -1.025464336 | 0.04505299 |
| AT1G04107 | 66.973443 | 1.018317248 | 0.04534628 |
| AT1G14750 | 36.29617011 | 1.292581239 | 0.046118809 |
| AT1G13970 | 32.38194463 | 1.3089261 | 0.04639387 |
| AT5G08150 | 62.03744472 | -1.10446322 | 0.046402246 |
| AT1G14260 | 44.94041593 | 1.183531455 | 0.046935744 |
| AT3G50300 | 11.11999676 | 2.236357304 | 0.046935744 |
| AT5G58570 | 40.12843706 | -1.221831901 | 0.047094695 |
| AT4G28500 | 10.8924526 | -2.43482181 | 0.047138161 |

| | | | |
|---|---|---|---|
| AT2G40750 | 20.55249396 | -1.733477739 | 0.047484276 |
| AT4G01895 | 40.35995563 | 1.346548645 | 0.047515961 |
| AT1G13540 | 54.18734637 | -1.008451812 | 0.047887689 |
| AT4G19820 | 35.69144754 | -1.324890341 | 0.048136566 |
| AT1G21120 | 729.3610668 | 1.095032645 | 0.048917301 |
| AT2G05235 | 18.86045595 | -1.714195624 | 0.048917301 |
| AT3G19620 | 14.24946795 | -2.320770182 | 0.049589251 |
| AT2G30750 | 1591.201879 | 1.446512689 | 0.049880794 |
| AT2G38750 | 67.05698265 | -1.09342765 | 0.049913371 |
| AT3G44550 | 40.99973517 | 1.321425486 | 0.049927285 |

**-m; *f* -m vs. *f* -K, *f* -m vs. *f* -Q and *f* -m vs. *f* -R along with the baseMean, log2-foldchange, adjusted p-value a**

| annotation |
|---|
| AT3G47340.ASN1.glutamine-dependent asparagine synthase 1.Gene |
| AT1G11260.STP1.sugar transporter 1.Gene |
| AT1G12240.ATBETAFRUCT4.Glycosyl hydrolases family 32 protein.Gene |
| AT4G35770.SEN1.Rhodanese/Cell cycle control phosphatase superfamily protein.Gene |
| AT1G03090.MCCA.methylcrotonyl-CoA carboxylase alpha chain.Gene |
| AT4G25580.AT4G25580.CAP160 protein.Gene |
| AT5G63160.BT1.BTB and TAZ domain protein 1.Gene |
| AT5G56870.BGAL4.beta-galactosidase 4.Gene |
| AT2G34430.LHB1B1.light-harvesting chlorophyll-protein complex II subunit B1.Gene |
| AT1G10070.BCAT-2.branched-chain amino acid transaminase 2.Gene |
| AT5G44120.CRA1.RmlC-like cupins superfamily protein.Gene |
| AT4G36850.AT4G36850.PQ-loop repeat family protein / transmembrane family protein.Gene |
| AT3G05727.AT3G05727.S locus-related glycoprotein 1 (SLR1) binding pollen coat protein family.Gene |
| AT5G41080.GDPD2.PLC-like phosphodiesterases superfamily protein.Gene |
| AT4G26260.MIOX4.myo-inositol oxygenase 4.Gene |
| AT1G75945.AT1G75945.hypothetical protein.Gene |
| AT3G16770.EBP.ethylene-responsive element binding protein.Gene |
| AT1G02660.AT1G02660.alpha/beta-Hydrolases superfamily protein.Gene |
| AT5G49360.BXL1.beta-xylosidase 1.Gene |
| AT1G21400.AT1G21400.Thiamin diphosphate-binding fold (THDP-binding) superfamily protein.Gene |
| AT5G23010.MAM1.methylthioalkylmalate synthase 1.Gene |
| AT1G08630.THA1.threonine aldolase 1.Gene |
| AT3G61198.AT3G61198..Gene |
| AT5G27920.AT5G27920.F-box family protein.Gene |
| AT3G15450.AT3G15450.aluminum induced protein with YGL and LRDR motifs.Gene |
| AT4G32870.AT4G32870.Polyketide cyclase/dehydrase and lipid transport superfamily protein.Gene |
| AT1G80160.GLYI7.Lactoylglutathione lyase / glyoxalase I family protein.Gene |
| AT3G06850.BCE2.2-oxoacid dehydrogenases acyltransferase family protein.Gene |
| AT5G09440.EXL4.EXORDIUM like 4.Gene |
| AT3G13450.DIN4.Transketolase family protein.Gene |
| AT5G07080.AT5G07080.HXXXD-type acyl-transferase family protein.Gene |
| AT5G18170.GDH1.glutamate dehydrogenase 1.Gene |
| AT1G77210.STP14.sugar transporter 14.Gene |
| AT5G51970.AT5G51970.GroES-like zinc-binding alcohol dehydrogenase family protein.Gene |
| AT3G45730.AT3G45730.hypothetical protein.Gene |
| AT1G10340.AT1G10340.Ankyrin repeat family protein.Gene |
| AT4G37870.PCK1.phosphoenolpyruvate carboxykinase 1.Gene |
| AT3G56360.AT3G56360.hypothetical protein.Gene |
| AT4G34030.MCCB.3-methylcrotonyl-CoA carboxylase.Gene |
| AT4G27450.AT4G27450.aluminum induced protein with YGL and LRDR motifs.Gene |
| AT3G19390.AT3G19390.Granulin repeat cysteine protease family protein.Gene |

AT3G00800..novel transcribed region.Gene
AT4G15530.PPDK.pyruvate orthophosphate dikinase.Gene
AT1G13300.HRS1.myb-like transcription factor family protein.Gene
AT1G19540.AT1G19540.NmrA-like negative transcriptional regulator family protein.Gene
AT2G33830.AT2G33830.Dormancy/auxin associated family protein.Gene
AT5G02020.SIS.E3 ubiquitin-protein ligase RLIM-like protein.Gene
AT2G38400.AGT3.alanine:glyoxylate aminotransferase 3.Gene
AT1G66760.AT1G66760.MATE efflux family protein.Gene
AT1G66180.AT1G66180.Eukaryotic aspartyl protease family protein.Gene
AT2G17880.AT2G17880.Chaperone DnaJ-domain superfamily protein.Gene
AT1G79360.OCT2.organic cation/carnitine transporter 2.Gene
AT1G15040.GAT1_2.1.Class I glutamine amidotransferase-like superfamily protein.Gene
AT1G19530.AT1G19530.DNA polymerase epsilon catalytic subunit A.Gene
AT3G61060.PP2-A13.phloem protein 2-A13.Gene
AT4G19160.AT4G19160.transglutaminase family protein.Gene
AT2G39980.AT2G39980.HXXXD-type acyl-transferase family protein.Gene
AT5G25980.TGG2.glucoside glucohydrolase 2.Gene
AT5G64110.AT5G64110.Peroxidase superfamily protein.Gene
AT1G79700.WRI4.Integrase-type DNA-binding superfamily protein.Gene
AT1G22500.ATL15.RING/U-box superfamily protein.Gene
AT1G06570.PDS1.4-hydroxyphenylpyruvate dioxygenase.Gene
AT5G21170.AKINBETA1.5'-AMP-activated protein kinase beta-2 subunit protein.Gene
AT5G13930.TT4.Chalcone and stilbene synthase family protein.Gene
AT5G24530.DMR6.2-oxoglutarate (2OG) and Fe(II)-dependent oxygenase superfamily protein.Gene
AT2G36120.DOT1.Glycine-rich protein family.Gene
AT3G29240.AT3G29240.PPR containing protein (DUF179).Gene
AT5G53970.TAT7.Tyrosine transaminase family protein.Gene
AT5G21020.AT5G21020.transmembrane protein.Gene
AT2G43620.AT2G43620.Chitinase family protein.Gene
AT4G28040.UMAMIT33.nodulin MtN21 /EamA-like transporter family protein.Gene
AT1G26770.EXPA10.expansin A10.Gene
AT3G20395.AT3G20395.RING/U-box superfamily protein.Gene
AT3G45300.IVD.isovaleryl-CoA-dehydrogenase.Gene
AT1G55920.SERAT2.1.serine acetyltransferase 2.1.Gene
AT5G44585.AT5G44585.hypothetical protein.Gene
AT2G36490.DML1.demeter-like 1.Gene
AT4G25630.FIB2.fibrillarin 2.Gene
AT5G39520.AT5G39520.hypothetical protein (DUF1997).Gene
AT4G39950.CYP79B2.cytochrome P450, family 79, subfamily B, polypeptide 2.Gene
AT3G20440.EMB2729.Alpha amylase family protein.Gene
AT3G58990.IPMI1.isopropylmalate isomerase 1.Gene
AT2G32150.AT2G32150.Haloacid dehalogenase-like hydrolase (HAD) superfamily protein.Gene
AT3G45290.MLO3.Seven transmembrane MLO family protein.Gene
AT5G09220.AAP2.amino acid permease 2.Gene
AT2G44380.AT2G44380.Cysteine/Histidine-rich C1 domain family protein.Gene
AT1G78370.GSTU20.glutathione S-transferase TAU 20.Gene
AT3G21870.CYCP2.1.cyclin p2.1.Gene

AT5G01740.AT5G01740.Nuclear transport factor 2 (NTF2) family protein.Gene
AT5G09530.PELPK1.hydroxyproline-rich glycoprotein family protein.Gene
AT2G43100.IPMI2.isopropylmalate isomerase 2.Gene
AT5G05250.AT5G05250.hypothetical protein.Gene
AT4G12500.AT4G12500.Bifunctional inhibitor/lipid-transfer protein/seed storage 2S albumin superfamily prc
AT5G25260.AT5G25260.SPFH/Band 7/PHB domain-containing membrane-associated protein family.Gene
AT4G21990.APR3.APS reductase 3.Gene
AT1G15010.AT1G15010.mediator of RNA polymerase II transcription subunit.Gene
AT1G76690.OPR2.12-oxophytodienoate reductase 2.Gene
AT4G28420.AT4G28420.Tyrosine transaminase family protein.Gene
AT5G52300.LTI65.CAP160 protein.Gene
AT3G48390.AT3G48390.MA3 domain-containing protein.Gene
AT2G18193.AT2G18193.P-loop containing nucleoside triphosphate hydrolases superfamily protein.Gene
AT4G33960.AT4G33960.hypothetical protein.Gene
AT1G66700.PXMT1.S-adenosyl-L-methionine-dependent methyltransferases superfamily protein.Gene
AT5G56100.AT5G56100.glycine-rich protein / oleosin.Gene
AT2G39450.MTP11.Cation efflux family protein.Gene
AT5G14200.IMD1.isopropylmalate dehydrogenase 1.Gene
AT3G26830.PAD3.Cytochrome P450 superfamily protein.Gene
AT1G02640.BXL2.beta-xylosidase 2.Gene
AT3G56400.WRKY70.WRKY DNA-binding protein 70.Gene
AT5G13170.SAG29.senescence-associated gene 29.Gene
AT2G47460.MYB12.myb domain protein 12.Gene
AT1G17170.GSTU24.glutathione S-transferase TAU 24.Gene
AT1G70830.MLP28.MLP-like protein 28.Gene
AT2G03865..novel transcribed region; detected in root, carpel, root apical meristem, receptacle, leaf, dark-gr
AT1G35670.CDPK2.calcium-dependent protein kinase 2.Gene
AT1G17745.PGDH.D-3-phosphoglycerate dehydrogenase.Gene
AT5G14180.MPL1.Myzus persicae-induced lipase 1.Gene
AT2G40420.AT2G40420.Transmembrane amino acid transporter family protein.Gene
AT3G54600.DJ1F.Class I glutamine amidotransferase-like superfamily protein.Gene
AT3G28930.AIG2.AIG2-like (avirulence induced gene) family protein.Gene
AT3G06355...Gene
AT3G60120.BGLU27.beta glucosidase 27.Gene
AT2G27380.EPR1.extensin proline-rich 1.Gene
AT3G02480.AT3G02480.Late embryogenesis abundant protein (LEA) family protein.Gene
AT1G62580.NOGC1.flavin containing monooxygenase FMO GS-OX-like protein.Gene
AT5G55970.AT5G55970.RING/U-box superfamily protein.Gene
AT5G62920.ARR6.response regulator 6.Gene
AT5G49480.CP1.Ca2 -binding protein 1.Gene
AT5G50915.AT5G50915.basic helix-loop-helix (bHLH) DNA-binding superfamily protein.Gene
AT3G19710.BCAT4.branched-chain aminotransferase4.Gene
AT3G16150.ASPGB1.N-terminal nucleophile aminohydrolases (Ntn hydrolases) superfamily protein.Gene
AT1G29395.COR413IM1.COLD REGULATED 314 INNER MEMBRANE 1.Gene
AT1G69252.AT1G69252..Gene
AT2G44460.BGLU28.beta glucosidase 28.Gene
AT2G39800.P5CS1.delta1-pyrroline-5-carboxylate synthase 1.Gene

AT1G52690.LEA7.Late embryogenesis abundant protein (LEA) family protein.Gene
AT1G16400.CYP79F2.cytochrome P450, family 79, subfamily F, polypeptide 2.Gene
AT2G19810.OZF1.CCCH-type zinc finger family protein.Gene
AT5G46730.AT5G46730.glycine-rich protein.Gene
AT5G52310.LTI78.low-temperature-responsive protein 78 (LTI78) / desiccation-responsive protein 29A (RD2
AT1G45201.TLL1.triacylglycerol lipase-like 1.Gene
AT5G15960.KIN1.stress-responsive protein (KIN1) / stress-induced protein (KIN1).Gene
AT3G54640.TSA1.tryptophan synthase alpha chain.Gene
AT2G41190.AT2G41190.Transmembrane amino acid transporter family protein.Gene
AT2G22470.AGP2.arabinogalactan protein 2.Gene
AT1G74930.ORA47.Integrase-type DNA-binding superfamily protein.Gene
AT1G16410.CYP79F1.cytochrome p450 79f1.Gene
AT3G02020.AK3.aspartate kinase 3.Gene
AT2G29490.GSTU1.glutathione S-transferase TAU 1.Gene
AT3G23880.AT3G23880.F-box and associated interaction domains-containing protein.Gene
AT4G13770.CYP83A1.cytochrome P450, family 83, subfamily A, polypeptide 1.Gene
AT1G33960.AIG1.P-loop containing nucleoside triphosphate hydrolases superfamily protein.Gene
AT2G04050.AT2G04050.MATE efflux family protein.Gene
AT2G44370.AT2G44370.Cysteine/Histidine-rich C1 domain family protein.Gene
AT5G14580.AT5G14580.polyribonucleotide nucleotidyltransferase.Gene
AT3G15460.AT3G15460.Ribosomal RNA processing Brix domain protein.Gene
AT1G03400.AT1G03400.2-oxoglutarate (2OG) and Fe(II)-dependent oxygenase superfamily protein.Gene
AT1G19610.PDF1.4.defensin-like protein.Gene
AT5G58860.CYP86A1.cytochrome P450, family 86, subfamily A, polypeptide 1.Gene
AT5G11420.AT5G11420.transmembrane protein, putative (Protein of unknown function, DUF642).Gene
AT1G10585.AT1G10585.basic helix-loop-helix (bHLH) DNA-binding superfamily protein.Gene
AT1G21110.IGMT3.O-methyltransferase family protein.Gene
AT3G25250.AGC2-1.AGC (cAMP-dependent, cGMP-dependent and protein kinase C) kinase family protein.G
AT2G40340.DREB2C.Integrase-type DNA-binding superfamily protein.Gene
AT5G65140.TPPJ.Haloacid dehalogenase-like hydrolase (HAD) superfamily protein.Gene
AT4G28703.AT4G28703.RmlC-like cupins superfamily protein.Gene
AT1G74710.EDS16.ADC synthase superfamily protein.Gene
AT5G57240.ORP4C.OSBP(oxysterol binding protein)-related protein 4C.Gene
AT1G26240.AT1G26240.Proline-rich extensin-like family protein.Gene
AT3G52720.ACA1.alpha carbonic anhydrase 1.Gene
AT5G64100.AT5G64100.Peroxidase superfamily protein.Gene
AT5G05965.AT5G05965.cell wall RBR3-like protein.Gene
AT3G16670.AT3G16670.Pollen Ole e 1 allergen and extensin family protein.Gene
AT1G13530.AT1G13530.hypothetical protein (DUF1262).Gene
AT2G21820.AT2G21820.seed maturation protein.Gene
AT1G74080.MYB122.myb domain protein 122.Gene
AT4G02845..novel transcribed region.Gene
AT5G14070.ROXY2.Thioredoxin superfamily protein.Gene
AT5G52900.MAKR6.membrane-associated kinase regulator.Gene
AT1G06957...Gene
AT3G59550.SYN3.Rad21/Rec8-like family protein.Gene
AT1G68500.AT1G68500.hypothetical protein.Gene

AT1G09420.G6PD4.glucose-6-phosphate dehydrogenase 4.Gene
AT1G51860.AT1G51860.Leucine-rich repeat protein kinase family protein.Gene
AT3G61890.HB-12.homeobox 12.Gene
AT3G62950.AT3G62950.Thioredoxin superfamily protein.Gene
AT5G13210.AT5G13210.Uncharacterized conserved protein UCP015417, vWA.Gene
AT5G26920.CBP60G.Cam-binding protein 60-like G.Gene
AT1G52000.AT1G52000.Mannose-binding lectin superfamily protein.Gene
AT2G25460.AT2G25460.EEIG1/EHBP1 protein amino-terminal domain protein.Gene
AT2G19670.PRMT1A.protein arginine methyltransferase 1A.Gene
AT4G34250.KCS16.3-ketoacyl-CoA synthase 16.Gene
AT1G67980.CCOAMT.caffeoyl-CoA 3-O-methyltransferase.Gene
AT2G41870.AT2G41870.Remorin family protein.Gene
AT1G05340.AT1G05340.cysteine-rich TM module stress tolerance protein.Gene
AT2G23540.AT2G23540.GDSL-like Lipase/Acylhydrolase superfamily protein.Gene
AT5G40990.GLIP1.GDSL lipase 1.Gene
AT1G21250.WAK1.cell wall-associated kinase.Gene
AT3G10110.MEE67.Mitochondrial import inner membrane translocase subunit Tim17/Tim22/Tim23 family p
AT5G10180.SULTR2.1.slufate transporter 2.1.Gene
AT2G48130.AT2G48130.Bifunctional inhibitor/lipid-transfer protein/seed storage 2S albumin superfamily prc
AT4G38080.AT4G38080.hydroxyproline-rich glycoprotein family protein.Gene
AT1G48130.PER1.1-cysteine peroxiredoxin 1.Gene
AT1G14950.AT1G14950.Polyketide cyclase/dehydrase and lipid transport superfamily protein.Gene
AT3G05955...Gene
AT5G54740.SESA5.seed storage albumin 5.Gene
AT5G25490.AT5G25490.Ran BP2/NZF zinc finger-like superfamily protein.Gene
AT3G06780.AT3G06780.glycine-rich protein.Gene
AT3G22840.ELIP1.Chlorophyll A-B binding family protein.Gene
AT3G05400.AT3G05400.Major facilitator superfamily protein.Gene
AT5G42530.AT5G42530.hypothetical protein.Gene
AT4G23280.CRK20.cysteine-rich RLK (RECEPTOR-like protein kinase) 20.Gene
AT4G28460.AT4G28460.transmembrane protein.Gene
AT1G17180.GSTU25.glutathione S-transferase TAU 25.Gene
AT5G66480.AT5G66480.bacteriophage N4 adsorption B protein.Gene
AT5G15120.AT5G15120.2-aminoethanethiol dioxygenase, putative (DUF1637).Gene
AT2G02990.RNS1.ribonuclease 1.Gene
AT5G44260.AT5G44260.Zinc finger C-x8-C-x5-C-x3-H type family protein.Gene
AT3G09405.AT3G09405.Pectinacetylesterase family protein.Gene
AT1G04800.AT1G04800.glycine-rich protein.Gene
AT4G08770.Prx37.Peroxidase superfamily protein.Gene
AT5G26230.MAKR1.membrane-associated kinase regulator.Gene
AT2G22510.AT2G22510.hydroxyproline-rich glycoprotein family protein.Gene
AT2G42900.AT2G42900.Plant basic secretory protein (BSP) family protein.Gene
AT5G66400.RAB18.Dehydrin family protein.Gene
AT1G78970.LUP1.lupeol synthase 1.Gene
AT5G02780.GSTL1.glutathione transferase lambda 1.Gene
AT1G74670.GASA6.Gibberellin-regulated family protein.Gene
AT4G27160.SESA3.seed storage albumin 3.Gene

AT1G56430.NAS4.nicotianamine synthase 4.Gene
AT4G23890.NdhS.NAD(P)H-quinone oxidoreductase subunit S.Gene
AT5G10946.AT5G10946.hypothetical protein.Gene
AT1G07620.ATOBGM.GTP-binding protein Obg/CgtA.Gene
AT1G17960.AT1G17960.Threonyl-tRNA synthetase.Gene
AT5G22580.AT5G22580.Stress responsive A/B Barrel Domain-containing protein.Gene
AT3G49410.AT3G49410.Transcription factor IIIC, subunit 5.Gene
AT5G03350.AT5G03350.Legume lectin family protein.Gene
AT4G18940.AT4G18940.RNA ligase/cyclic nucleotide phosphodiesterase family protein.Gene
AT2G14247.AT2G14247.Expressed protein.Gene
AT4G27150.SESA2.seed storage albumin 2.Gene
AT2G47770.TSPO.TSPO(outer membrane tryptophan-rich sensory protein)-like protein.Gene
AT3G47780.ABCA7.ABC2 homolog 6.Gene
AT2G05695...Gene
AT5G17350.AT5G17350.hypothetical protein.Gene
AT4G20970.AT4G20970.basic helix-loop-helix (bHLH) DNA-binding superfamily protein.Gene
AT1G67865.AT1G67865.hypothetical protein.Gene
AT5G65600.AT5G65600.Concanavalin A-like lectin protein kinase family protein.Gene
AT1G55990.AT1G55990.glycine-rich protein.Gene
AT1G19960.AT1G19960.transcription factor.Gene
AT2G32870.AT2G32870.TRAF-like family protein.Gene
AT1G16150.WAKL4.wall associated kinase-like 4.Gene
AT1G53490.HEI10.RING/U-box superfamily protein.Gene
AT2G14160.AT2G14160.RNA-binding (RRM/RBD/RNP motifs) family protein.Gene
AT2G35380.AT2G35380.Peroxidase superfamily protein.Gene
AT5G37690.AT5G37690.SGNH hydrolase-type esterase superfamily protein.Gene
AT4G20390.AT4G20390.Uncharacterized protein family (UPF0497).Gene
AT1G14880.PCR1.PLANT CADMIUM RESISTANCE 1.Gene
AT2G01422.AT2G01422..Gene
AT2G41280.M10.late embryogenesis abundant protein (M10) / LEA protein M10.Gene
AT1G31580.ECS1.ECS1.Gene
AT5G06530.ABCG22.ABC-2 type transporter family protein.Gene
AT3G18400.NAC058.NAC domain containing protein 58.Gene
AT1G61440.AT1G61440.S-locus lectin protein kinase family protein.Gene
AT5G58390.AT5G58390.Peroxidase superfamily protein.Gene
AT4G01450.UMAMIT30.nodulin MtN21 /EamA-like transporter family protein.Gene
AT5G22200.AT5G22200.Late embryogenesis abundant (LEA) hydroxyproline-rich glycoprotein family.Gene
AT1G01110.IQD18.IQ-domain 18.Gene
AT2G40370.LAC5.laccase 5.Gene
AT3G05770.AT3G05770.hypothetical protein.Gene
AT2G27690.CYP94C1.cytochrome P450, family 94, subfamily C, polypeptide 1.Gene
AT3G12710.AT3G12710.DNA glycosylase superfamily protein.Gene
AT1G51140.FBH3.basic helix-loop-helix (bHLH) DNA-binding superfamily protein.Gene
AT1G18710.MYB47.myb domain protein 47.Gene
AT5G38940.AT5G38940.RmlC-like cupins superfamily protein.Gene
AT1G69920.GSTU12.glutathione S-transferase TAU 12.Gene
AT4G12030.BAT5.bile acid transporter 5.Gene

AT5G05960.AT5G05960.Bifunctional inhibitor/lipid-transfer protein/seed storage 2S albumin superfamily pro
AT1G52450.AT1G52450.Ubiquitin carboxyl-terminal hydrolase-related protein.Gene
AT3G50440.MES10.methylesterase.Gene
AT4G16000.AT4G16000.hypothetical protein.Gene
AT2G26010.PDF1.3.plant defensin 1.3.Gene
AT5G02890.AT5G02890.HXXXD-type acyl-transferase family protein.Gene
AT1G20850.XCP2.xylem cysteine peptidase 2.Gene
AT1G64370.AT1G64370.filaggrin-like protein.Gene
AT1G51380.AT1G51380.DEA(D/H)-box RNA helicase family protein.Gene
AT1G67910.AT1G67910.hypothetical protein.Gene
AT4G37290.AT4G37290.transmembrane protein.Gene
AT1G60750.AT1G60750.NAD(P)-linked oxidoreductase superfamily protein.Gene
AT5G38030.AT5G38030.MATE efflux family protein.Gene
AT1G60110.AT1G60110.Mannose-binding lectin superfamily protein.Gene
AT5G38710.AT5G38710.Methylenetetrahydrofolate reductase family protein.Gene
AT3G16120.AT3G16120.Dynein light chain type 1 family protein.Gene
AT2G26695.AT2G26695.Ran BP2/NZF zinc finger-like superfamily protein.Gene
AT1G15050.IAA34.indole-3-acetic acid inducible 34.Gene
AT5G09580.AT5G09580.heat shock protein.Gene
AT4G27460.CBSX5.Cystathionine beta-synthase (CBS) family protein.Gene
AT3G49580.LSU1.response to low sulfur 1.Gene
AT4G17030.EXLB1.expansin-like B1.Gene
AT3G14580.AT3G14580.Pentatricopeptide repeat (PPR) superfamily protein.Gene
AT5G19600.SULTR3.5.sulfate transporter 3.5.Gene
AT1G68850.AT1G68850.Peroxidase superfamily protein.Gene
AT3G06520.AT3G06520.agenet domain-containing protein.Gene
AT3G28580.AT3G28580.P-loop containing nucleoside triphosphate hydrolases superfamily protein.Gene
AT3G22740.HMT3.homocysteine S-methyltransferase 3.Gene
AT5G06930.AT5G06930.nucleolar-like protein.Gene
AT1G16850.AT1G16850.transmembrane protein.Gene
AT1G07400.AT1G07400.HSP20-like chaperones superfamily protein.Gene
AT5G03615...Gene
AT5G60730.AT5G60730.Anion-transporting ATPase.Gene
AT3G21370.BGLU19.beta glucosidase 19.Gene
AT3G50800.AT3G50800.hypothetical protein.Gene
AT4G33610.AT4G33610.glycine-rich protein.Gene
AT5G57550.XTH25.xyloglucan endotransglucosylase/hydrolase 25.Gene
AT1G34670.MYB93.myb domain protein 93.Gene
AT3G25882.NIMIN-2.NIM1-interacting 2.Gene
AT4G23150.CRK7.cysteine-rich RLK (RECEPTOR-like protein kinase) 7.Gene
AT1G76960.AT1G76960.transmembrane protein.Gene
AT5G48900.AT5G48900.Pectin lyase-like superfamily protein.Gene
AT1G19510.RL5.RAD-like 5.Gene
AT4G21650.AT4G21650.Subtilase family protein.Gene
AT5G59320.LTP3.lipid transfer protein 3.Gene
AT5G12030.HSP17.6A.heat shock protein 17.6A.Gene
AT2G17036.AT2G17036.F-box SKIP23-like protein (DUF295).Gene

AT2G05895...Gene
AT5G13580.ABCG6.ABC-2 type transporter family protein.Gene
AT4G22070.WRKY31.WRKY DNA-binding protein 31.Gene
AT4G30460.AT4G30460.glycine-rich protein.Gene
AT3G55710.AT3G55710.UDP-Glycosyltransferase superfamily protein.Gene
AT1G30720.AT1G30720.FAD-binding Berberine family protein.Gene
AT3G53530.NAKR3.Chloroplast-targeted copper chaperone protein.Gene
AT5G35580.AT5G35580.Protein kinase superfamily protein.Gene
AT2G45050.GATA2.GATA transcription factor 2.Gene
AT1G64360.AT1G64360.hypothetical protein.Gene
AT1G13609.AT1G13609.Defensin-like (DEFL) family protein.Gene
AT5G04200.MC9.metacaspase 9.Gene
AT1G27670.AT1G27670.transmembrane protein.Gene
AT5G37990.AT5G37990.S-adenosyl-L-methionine-dependent methyltransferase superfamily protein.Gene
AT5G42800.DFR.dihydroflavonol 4-reductase.Gene
AT4G04223.AT4G04223..Gene
AT5G62280.AT5G62280.DUF1442 family protein (DUF1442).Gene
AT1G64940.CYP89A6.cytochrome P450, family 87, subfamily A, polypeptide 6.Gene
AT2G40010.AT2G40010.Ribosomal protein L10 family protein.Gene
AT5G15970.KIN2.stress-responsive protein (KIN2) / stress-induced protein (KIN2) / cold-responsive protein (C
AT2G43590.AT2G43590.Chitinase family protein.Gene
AT1G74460.AT1G74460.GDSL-like Lipase/Acylhydrolase superfamily protein.Gene
AT1G67750.AT1G67750.Pectate lyase family protein.Gene
AT3G21550.DMP2.transmembrane protein, putative (DUF679 domain membrane protein 2).Gene
AT2G43000.NAC042.NAC domain containing protein 42.Gene
AT3G16360.AHP4.HPT phosphotransmitter 4.Gene
AT1G22160.AT1G22160.senescence-associated family protein (DUF581).Gene
AT2G34360.AT2G34360.MATE efflux family protein.Gene
AT1G60590.AT1G60590.Pectin lyase-like superfamily protein.Gene
AT2G16890.AT2G16890.UDP-Glycosyltransferase superfamily protein.Gene
AT4G27140.SESA1.seed storage albumin 1.Gene
AT3G11430.GPAT5.glycerol-3-phosphate acyltransferase 5.Gene
AT1G66040.VIM4.Zinc finger (C3HC4-type RING finger) family protein.Gene
AT1G15830.AT1G15830.hypothetical protein.Gene
AT3G01345.AT3G01345.Expressed protein.Gene
AT3G48340.CEP2.Cysteine proteinases superfamily protein.Gene
AT2G48140.EDA4.Bifunctional inhibitor/lipid-transfer protein/seed storage 2S albumin superfamily protein.G
AT3G01190.AT3G01190.Peroxidase superfamily protein.Gene
AT5G51810.GA20OX2.gibberellin 20 oxidase 2.Gene
AT3G54940.AT3G54940.Papain family cysteine protease.Gene
AT2G42870.PAR1.phy rapidly regulated 1.Gene
AT2G25680.MOT1.molybdate transporter 1.Gene
AT1G72870.AT1G72870.Disease resistance protein (TIR-NBS class).Gene
AT5G60760.AT5G60760.P-loop containing nucleoside triphosphate hydrolases superfamily protein.Gene
AT4G30450.AT4G30450.glycine-rich protein.Gene
AT5G35660.AT5G35660.Glycine-rich protein family.Gene
AT1G49960.AT1G49960.Xanthine/uracil permease family protein.Gene

AT2G23270.AT2G23270.transmembrane protein.Gene
AT2G32210.AT2G32210.cysteine-rich/transmembrane domain A-like protein.Gene
AT5G57560.TCH4.Xyloglucan endotransglucosylase/hydrolase family protein.Gene
AT3G21380.AT3G21380.Mannose-binding lectin superfamily protein.Gene
AT2G30370.CHAL.allergen-like protein.Gene
AT4G04450.WRKY42.WRKY family transcription factor.Gene
AT5G24330.ATXR6.TRITHORAX-RELATED PROTEIN 6.Gene
AT1G26380.AT1G26380.FAD-binding Berberine family protein.Gene
AT1G17285.AT1G17285.transmembrane protein.Gene
AT3G10870.MES17.methyl esterase 17.Gene
AT3G59480.AT3G59480.pfkB-like carbohydrate kinase family protein.Gene
AT5G18430.AT5G18430.GDSL-like Lipase/Acylhydrolase superfamily protein.Gene
AT2G32200.AT2G32200.cysteine-rich/transmembrane domain A-like protein.Gene
AT1G18320.AT1G18320.Mitochondrial import inner membrane translocase subunit Tim17/Tim22/Tim23 fam
AT5G43580.UPI.Serine protease inhibitor, potato inhibitor I-type family protein.Gene
AT1G74810.BOR5.HCO3- transporter family.Gene
AT1G11850.AT1G11850.transmembrane protein.Gene
AT5G07475.AT5G07475.Cupredoxin superfamily protein.Gene
AT4G28520.CRU3.cruciferin 3.Gene
AT2G00340..novel transcribed region.Gene
AT3G04320.AT3G04320.Kunitz family trypsin and protease inhibitor protein.Gene
AT5G21150.AGO9.Argonaute family protein.Gene
AT3G50760.GATL2.galacturonosyltransferase-like 2.Gene
AT4G22160.AT4G22160.hypothetical protein.Gene
AT5G23190.CYP86B1.cytochrome P450, family 86, subfamily B, polypeptide 1.Gene
AT3G15357.AT3G15357.phosphopantothenoylcysteine decarboxylase subunit.Gene
AT1G03100.AT1G03100.Pentatricopeptide repeat (PPR) superfamily protein.Gene
AT2G22590.AT2G22590.UDP-Glycosyltransferase superfamily protein.Gene
AT1G10990.AT1G10990.transmembrane protein.Gene
AT1G09867...Gene
AT5G23700.AT5G23700.coiled-coil protein.Gene
AT3G18170.AT3G18170.Glycosyltransferase family 61 protein.Gene
AT4G23990.CSLG3.cellulose synthase like G3.Gene
AT5G15500.AT5G15500.Ankyrin repeat family protein.Gene
AT3G16530.AT3G16530.Legume lectin family protein.Gene
AT5G16410.AT5G16410.HXXXD-type acyl-transferase family protein.Gene
AT4G33070.AT4G33070.Thiamine pyrophosphate dependent pyruvate decarboxylase family protein.Gene
AT1G63310.AT1G63310.hypothetical protein.Gene
AT5G10120.AT5G10120.Ethylene insensitive 3 family protein.Gene
AT3G14440.NCED3.nine-cis-epoxycarotenoid dioxygenase 3.Gene
AT1G09475..novel transcribed region.Gene
AT4G30250.AT4G30250.P-loop containing nucleoside triphosphate hydrolases superfamily protein.Gene
AT2G46640.AT2G46640.NAD-dependent protein deacetylase HST1-like protein.Gene
AT5G43890.YUC5.Flavin-binding monooxygenase family protein.Gene
AT5G17040.AT5G17040.UDP-Glycosyltransferase superfamily protein.Gene
AT5G42210.AT5G42210.Major facilitator superfamily protein.Gene
AT3G09190.AT3G09190.Concanavalin A-like lectin family protein.Gene

AT1G02380.AT1G02380.transmembrane protein.Gene
AT3G55990.ESK1.trichome birefringence-like protein (DUF828).Gene
AT1G13800.FAC19.Tetratricopeptide repeat (TPR)-like superfamily protein.Gene
AT5G43180.AT5G43180.transmembrane protein, putative (Protein of unknown function, DUF599).Gene
AT1G51920.AT1G51920.transmembrane protein.Gene
AT5G24940.AT5G24940.Protein phosphatase 2C family protein.Gene
AT1G06553...Gene
AT4G15320.CSLB06.cellulose synthase-like B6.Gene
AT1G47980.AT1G47980.desiccation-like protein.Gene
AT4G26120.AT4G26120.Ankyrin repeat family protein / BTB/POZ domain-containing protein.Gene
AT3G49142.AT3G49142.Tetratricopeptide repeat (TPR)-like superfamily protein.Gene
AT3G59710.AT3G59710.NAD(P)-binding Rossmann-fold superfamily protein.Gene
AT3G00190..novel transcribed region.Gene
AT1G62540.FMO GS-OX2.flavin-monooxygenase glucosinolate S-oxygenase 2.Gene
AT3G00390..novel transcribed region.Gene
AT4G26990.AT4G26990.polyadenylate-binding protein interacting protein.Gene
AT4G19870.AT4G19870.Galactose oxidase/kelch repeat superfamily protein.Gene
AT5G61250.GUS1.glucuronidase 1.Gene
AT2G19970.AT2G19970.CAP (Cysteine-rich secretory proteins, Antigen 5, and Pathogenesis-related 1 protein
AT5G04150.BHLH101.basic helix-loop-helix (bHLH) DNA-binding superfamily protein.Gene
AT4G14120.AT4G14120.hypothetical protein.Gene
AT2G26290.ARSK1.root-specific kinase 1.Gene
AT1G30730.AT1G30730.FAD-binding Berberine family protein.Gene
AT4G13690.AT4G13690.RNA-binding protein.Gene
AT4G23160.CRK8.cysteine-rich RECEPTOR-like kinase.Gene
AT4G26960.AT4G26960.hypothetical protein.Gene
AT2G42540.COR15A.cold-regulated 15a.Gene
AT1G01680.PUB54.plant U-box 54.Gene
AT3G50400.AT3G50400.GDSL-like Lipase/Acylhydrolase superfamily protein.Gene
AT1G74490.AT1G74490.Protein kinase superfamily protein.Gene
AT4G21020.AT4G21020.Late embryogenesis abundant protein (LEA) family protein.Gene
AT3G45130.LAS1.lanosterol synthase 1.Gene
AT5G44582.AT5G44582.hypothetical protein.Gene
AT4G28720.YUC8.Flavin-binding monooxygenase family protein.Gene
AT1G02335.GL22.germin-like protein subfamily 2 member 2 precursor.Gene
AT3G00850..novel transcribed region.Gene
AT3G09480.AT3G09480.Histone superfamily protein.Gene
AT5G64870.AT5G64870.SPFH/Band 7/PHB domain-containing membrane-associated protein family.Gene
AT3G51540.AT3G51540.mucin-5AC-like protein.Gene
AT1G04107..novel transcribed region; detected in root, carpel, root apical meristem, receptacle, leaf, dark-gr
AT1G14750.SDS.Cyclin family protein.Gene
AT1G13970.AT1G13970.beta-hexosaminidase (DUF1336).Gene
AT5G08150.SOB5.suppressor of phytochrome b 5.Gene
AT1G14260.AT1G14260.RING/FYVE/PHD zinc finger superfamily protein.Gene
AT3G50300.AT3G50300.HXXXD-type acyl-transferase family protein.Gene
AT5G58570.AT5G58570.transmembrane protein.Gene
AT4G28500.NAC073.NAC domain containing protein 73.Gene

AT2G40750.WRKY54.WRKY DNA-binding protein 54.Gene
AT4G01895.AT4G01895.systemic acquired resistance (SAR) regulator protein NIMIN-1-like protein.Gene
AT1G13540.AT1G13540.hypothetical protein (DUF1262).Gene
AT4G19820.AT4G19820.Glycosyl hydrolase family protein with chitinase insertion domain-containing protein
AT1G21120.IGMT2.O-methyltransferase family protein.Gene
AT2G05235..novel transcribed region; detected in root, carpel, root apical meristem, receptacle, leaf, dark-gr
AT3G19620.AT3G19620.Glycosyl hydrolase family protein.Gene
AT2G30750.CYP71A12.cytochrome P450 family 71 polypeptide.Gene
AT2G38750.ANNAT4.annexin 4.Gene
AT3G44550.FAR5.fatty acid reductase 5.Gene

nd their annotation.

rown seedling, light-grown seedling, flower, pollen, aerial, shoot apical meristem.Gene

rown seedling, light-grown seedling, flower, aerial, shoot apical meristem.Gene
